# S_1_ basic leucine zipper transcription factors shape plant architecture by controlling C/N partitioning to apical and lateral organs

**DOI:** 10.1101/2023.05.23.541964

**Authors:** Philipp Kreisz, Alicia M. Hellens, Christian Fröschel, Markus Krischke, Daniel Maag, Regina Feil, Theresa Wildenhain, Jan Draken, Gabriel Braune, Leon Erdelitsch, Laura Cecchino, Tobias C. Wagner, Martin J. Mueller, Dirk Becker, John E. Lunn, Johannes Hanson, Christine A. Beveridge, Franziska Fichtner, Francois F. Barbier, Christoph Weiste

## Abstract

Plants exhibit an immense plasticity in their architecture. While the impact of hormonal regulation is well-characterised, the importance of sugar-signalling has just recently emerged. Here, we addressed which sugar-signalling components mediate the trade-off between growth of apical versus lateral meristems and how they control organ sink-strength. Thereby, we unravelled a novel developmental function of the sugar-controlled S_1_ basic-leucine-zipper (S_1_-bZIP) transcription factors in establishing global source-sink interactions. Applying comprehensive molecular, analytical, and genetic approaches, we demonstrate that S_1_-bZIPs operate in a redundant manner to control tissue-specific expression of defined *SWEET* sugar-transporters and the *GAT1_2.1* glutaminase. By these means, S_1_-bZIPs control carbohydrate (C)-channelling from source leaves to apical shoot and root organs and tune systemic organic nitrogen (N)-supply to restrict lateral organ formation by C/N depletion. Knowledge of the underlying mechanisms controlling plant C/N partitioning is of pivotal importance for breeding strategies to generate plants with desired architectural and nutritional characteristics.

## Introduction

Plants have an enormous capacity to modulate their architecture. However, many agronomically important plant species, as well as the model plant Arabidopsis (*Arabidopsis thaliana*) exhibit a growth dominance of the apical meristems, while the pre-established lateral shoot (axillary branch) and root (lateral root, LR) primordia remain initially dormant^1, 2^. This control of nutrient-intensive growth of lateral organs in support of apical meristems, referred to as apical dominance, is beneficial for the plant’s Darwinian fitness^3^. It prevents plants from being overgrown by neighbouring plants competing for light, water and minerals and ensures species conservation in fluctuating environments by concentrating limited resources on the primary inflorescence to allow seed maturation. As such, apical dominance provides a gateway for successful dispersal into competitive and resource-scarce habitats without the need for genetic change and is therefore a key determinant of plant evolutionary success.

Nevertheless, upon silique ripening and the concomitant decrease in carbohydrate (C) demand of the growing shoot tip, repression of axillary shoot outgrowth is released, allowing an upscaling of seed production^4^. Consistently, decapitation of the primary shoot apex, which results in rapid C reallocation to axillary branches^5^ or cultivation under high-light conditions that increase total C availability^6^ strongly reduce apical dominance. Similar observations have been made for LR growth, which is induced by either root apical meristem excision^7^, root sugar feeding or high-light cultivation^8, 9^, suggesting a coordinated, C-responsive expansion of shoot and root tissues^2, 9^. Indeed, recent studies demonstrate that C redistribution to lateral organs represents the initial trigger for primordia outgrowth and is dominant over the repressive effects of the well-established branching-control hormones auxin and strigolactone^5, 10^. Despite its pivotal importance, the underlying mechanisms that enable plants to integrate nutritional inputs and direct resources to apical or lateral meristems to modulate overall architecture remains largely unresolved.

However, major advances in uncovering transporter activities that facilitate C fluxes from the site of production (source) to consuming tissues (sink), assist in revealing the coordination of C partitioning. For instance, sugar fluxes in Arabidopsis originate largely from photosynthetically active rosette leaves, which act as strong C source tissue. While a significant fraction of leaf C is stored as starch, sucrose can passively diffuse from the leaf mesophyll cells either across plasmodesmata to neighbouring tissues or preferentially along the gradient to the apoplastic space via sucrose efflux transporter proteins (Sugars Will Eventually be Exported Transporters, SWEETs)^11^. For long-distance transport to distal sinks, incoming apoplastic sugars are actively loaded into the phloem companion cells by sucrose-H^+^ symporters of the SUCROSE TRANSPORTER family (SUC/SUTs) and are either unloaded symplastically or by SWEETs at the growing, C-demanding apical shoot, or root meristems^12^. This creates a constant flow of sugars from the source tissue to the distal apical meristems, while the proximal lateral sinks remain C-depleted, thereby maintaining apical growth dominance.

In addition to C supply, the availability of organic N is essential for meristem growth^4^. Most N uptake takes place in roots that absorb nitrate or ammonium via specialised membrane-localised transporters from the surrounding soil^13^. While ammonium serves directly as a substrate for root GLUTAMINE SYNTHETASE (GS) isozymes to form glutamine (Gln) from glutamate, nitrate is primarily transported through the xylem to the leaves, where it also feeds the Gln pool after initial enzymatic reduction to ammonium and final conversion by leaf GS^13^. Gln, the most abundant amino acid (AA) in plants, is used as a major transport form of organic N and acts as a central nitrogen donor for the biosynthesis of proteinogenic AAs or nucleotides for DNA replication. To control homeostasis of this key AA, Gln can be deaminated by glutaminases such as the recently identified GLUTAMINE AMIDO-TRANSFERASE 1_2.1 (GAT1_2.1) to initiate its degradation via the tricarboxylic acid cycle^14^. Importantly, loss-of-function mutant analysis has implicated *GAT1_2.1* in shoot branching control^15^.

In plants, two highly conserved and counteracting metabolic kinases, namely TARGET OF RAPAMYCIN (TOR) and SUCROSE-NON-FERMENTING1 RELATED KINASE1 (SnRK1), are known to be central components of a C/N status integrating system that controls plant growth and metabolism by reprogramming global gene expression and enzymatic activities^16–18^. Whereas TOR is activated by a high C/N ratio and promotes anabolism and plant growth responses, SnRK1 signalling is activated under energy-deficient conditions and induces catabolic processes and growth repression^17, 18^. Although functionally well-characterised in managing severe energy deprivation, recent work highlights the impact of SnRK1 and its catalytic subunit SnRK1α1 in controlling sink tissue development, such as seedling establishment^19^ and lateral root growth^20^, respectively. Although SnRK1 addresses multiple targets in this respect, the downstream group C basic leucine zipper (bZIP) transcription factor (TF) bZIP63, which preferentially heterodimerises with group S_1_-bZIPs upon phosphorylation^21, 22^, appears to implement key aspects of the SnRK1α1-driven transcriptional adaptation^19, 20, 23^. Consistently, SnRK1α1 strongly boosts the transactivation potential of group S_1_-bZIPs^24^, leading to the concept of an interconnected SnRK1-C/S_1_ bZIP heterodimerisation network^25^.

Along this line recent work highlights the importance of individual members of the highly homologous S_1_-bZIP TFs: *bZIP1*, *bZIP2*, *bZIP11*, *bZIP44* and *bZIP53* in mediating growth repression and metabolic adaptation^23, 26, 27^. As they possess an additional layer of post-transcriptional regulation through conserved uORFs in their 5’ leader sequences that confer sucrose-induced repression of translation, it is postulated that they have the intrinsic capacity to integrate sucrose availability into growth responses^28^. Although S_1_-bZIPs exhibit highly overlapping expression domains in source (leaf petioles of mature leaves) and sink (young leaves, root tip, flowers, and rosette core with branches)^29^ tissues, their functional implications have not been addressed, particularly due to limited availability of mutants and potential functional redundancy.

In this study, we focused our attention on unravelling the role of group S_1_-bZIP TFs in controlling energy-intensive lateral organ formation. Therefore, we used CRISPR/Cas9 to generate single and higher-order mutant combinations of S_1_-bZIPs and analysed their growth throughout the vegetative and reproductive phases. In contrast to wild-type (WT) plants, which showed a marked dominance of apical shoot and root growth, we observed a clear promotion of lateral organ growth at the expense of apical organs in higher-order mutants of S_1_-bZIPs, suggesting their importance for establishing apical dominance. Strikingly, removal of excessive C-demanding branches or increasing root C supply by sugar feeding rescued the impaired apical growth responses of the mutants. Therefore, we hypothesised that S_1_-bZIPs are required for C-channelling to apical meristems. Consistent with this hypothesis, mutants showed a strong reduction in sugar availability in the leaf apoplast and distal sinks, but an increase in proximal sinks (axillary branches) and leaf cells, indicative of an impaired sugar export from the latter. Further metabolic analyses revealed an additional function of S_1_-bZIPs in controlling root and shoot N (especially Gln) homeostasis. Applying comprehensive transcriptomics, molecular biology approaches and mutant analysis we identified specific clade 3 *SWEETs* and the glutaminase *GAT1_2.1* as direct S_1_ targets involved in controlling C and N supply. Therefore, we propose that the energy-controlled S_1_-bZIP TFs act as key regulators of shoot and root apical dominance by determining the C/N status of apical and lateral meristems.

## Results

### Higher-order mutants of group S_1_-bZIP TFs exhibit a growth repression of apical and growth promotion of lateral organs

In recent years, gain-of-function approaches implicated the low-energy responsive group S_1_-bZIP TFs in mediating systemic growth repression^26, 30^. Nevertheless, they were found to be expressed in specific C-sink and –source tissues under non-starved conditions, suggesting their involvement in sink-source communication and control of plant development^29^. However, a lack of mutants and potential functional redundancy among the highly homologous S_1_ members severely hampered their functional characterisation. To assess the impact of group S_1_ bZIP members in this respect, we used CRISPR/Cas9^31^ to generate individual single bZIP knockouts and higher-order mutant combinations, ranging from double to a full group S_1_ quintuple mutant (Supplementary Table 1). While single and double mutants showed no obvious alterations in vegetative growth (Supplementary Fig. 1a), a triple mutant of *bzip2*, *bzip11* and *bzip44* (*bzip2/11/44*) or a full group S_1_ quintuple mutant (*bzip1/2/11/44/53*) showed, after a WT-like expansion of the first true leaves, a slight (*bzip2/11/44*) to marked (*bzip1/2/11/44/53*) shoot growth retardation, respectively (Fig. 1a-b). This mutant phenotype was preceded by an impaired primary root growth response (Fig. 1c-d) and, surprisingly, an early lateral root emergence compared to WT (Fig. 1e).

**Fig. 1.**
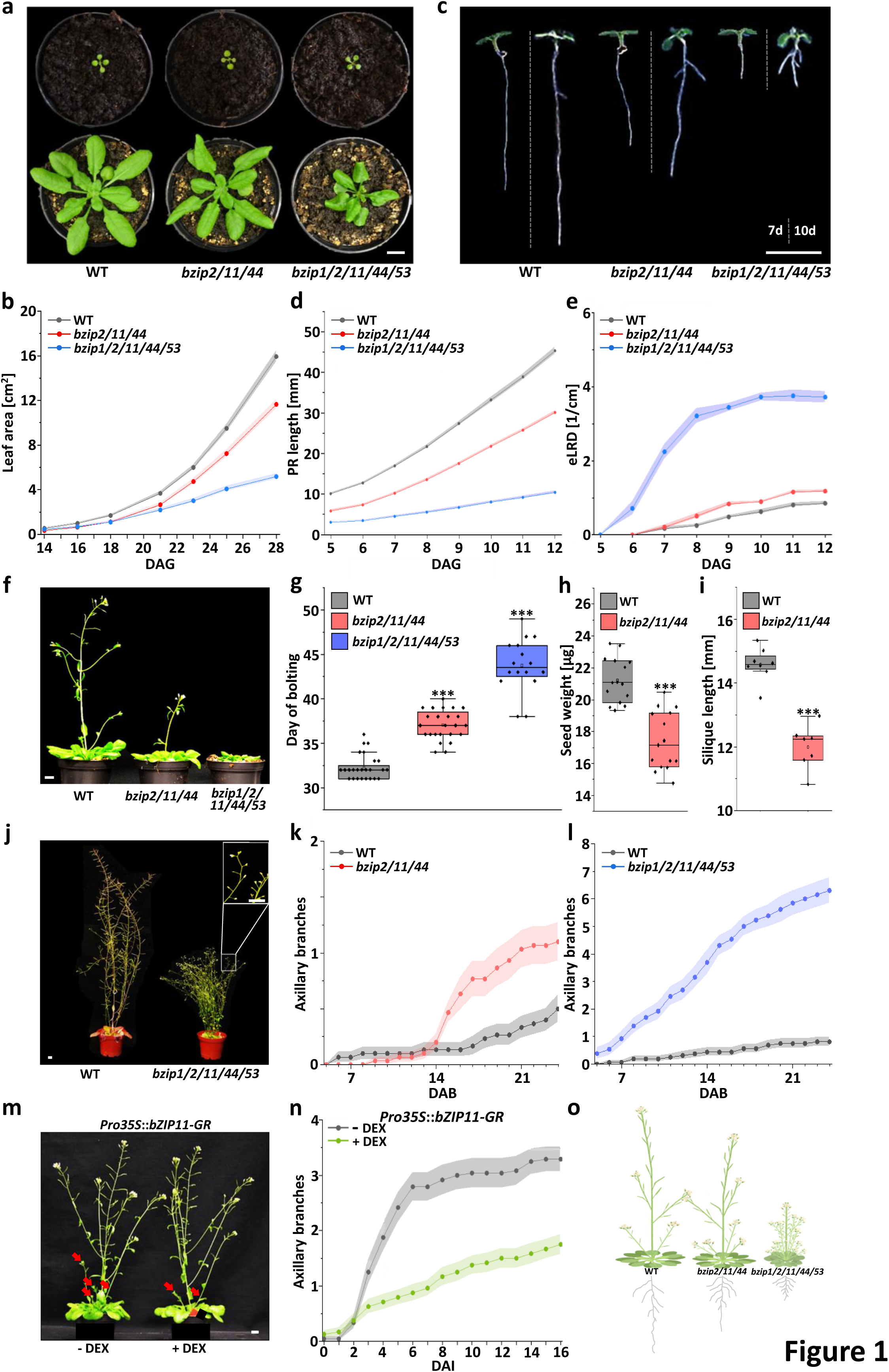
I Mutants of S_1_ bZIP TFs exhibit a growth repression of apical and growth promotion of lateral organs. **a**, Shoot growth of S_1_ bZIP mutants is impaired at late vegetative stage. Representative pictures of soil-grown 2-(upper row) and 4-(lower row) week-old WT, bZIP triple (*bzip2/-11/-44*) and quintuple (full group S_1,_ *bzip1/-2/-11/-44/-53}* mutants. Scale bar: 1cm. **b**, Mean leaf area (± SEM, shaded area, n=24 per genotype) of WT (black line), triple (red line) and quintuple (blue line) bZIP mutants from 14 to 28 days after germination (DAG). **c**, Primary root growth of S_1_ bZIP mutants is impaired. Representative picture of 7-(left) and 10-day-old (right) WT, bZIP triple and quintuple mutant seedlings. Scale bar: 1 cm. **d**, Mean PR length (± SEM, n=6, each replicate consisting of 10 plants per genotype) of WT (black line), bZIP triple (red line) and quintuple (blue line) seedlings was measured from 5 to 12 DAG. **e**, Emerged lateral root density (eLRD) over time of WT (black line) bZIP triple (red line) and quintuple (blue line) mutants. Given are mean values (± SEM, n=6, each replicate consisting of 10 plants per genotype). **f**, Bolting of S_1_ bZIP mutants is delayed. Representative picture of 6-week-old WT, bZIP triple and quintuple mutants. Scale bar: 1cm. **g**, Time of bolting of WT (black box), bZIP triple (red box) and quintuple (blue box) mutants. Given are mean values (n=16-24 per genotype). **h**, Mean seed weight (n=15 per genotype) of WT (black box) and bZIP triple (red box) mutants. **i**, Mean silique length (n=7-8 per genotype) of siliques 6 to 15 (counted from the inflorescence base) from 8-week-old WT (black box) and bZIP triple (red box) mutants. **j**, S_1_ bZIP mutants exhibit a high branching phenotype. Representative picture of 12-week-old WT and bZIP quintuple mutants. Scale bar: 1 cm Magnification shows high number of extremely short and frequently aborted siliques. Scale bar: 1 cm. **k**, Mean number (± SEM, n=30 per genotype) of axillary branches of WT (black line) and bZIP triple (red line) or (**l**) WT (black line) and bZIP quintuple mutants (blue line) (± SEM, n=13-16 per genotype). Outgrown axillary branches (g1cm) were counted starting 5 days after bolting (DAB). **m**, Local overexpression of bZIP11 in rosette cores represses axillary branching but does not affect primary shoot growth (**Supplementary** Fig. 1k). Representative picture of mock treated (solvent without dexamethasone, – DEX) and DEX induced (+ DEX) Pro35S::*bZ/P11*-GR plants at 7 days after start of induction (DAI). Treatment started prior to first axillary branching when primary inflorescences reached ∼10 cm. DEX or the solvent control was applied every second day to the rosette cores. Red arrows mark outgrown axillary branches. Scale bar: 1cm. **n**, Mean number (± SEM, n=24 per treatment) of outgrown axillary branches of Pro35S::*bZ/P11*-GR plants after recurring mock (black line) or DEX (green line) treatment. **o**, Schematic picture of differences in apical and lateral organ growth of WT and S_1_ bZIP mutants. Plants were either grown on soil (**Fig. 1a, b, f-n**) or on ½-strength MS medium (**Fig. 1c-e**) under long day conditions at 100 µmol photons m^-2^ s^-1^. Statistically significant differences were determined by Student’s *t*-test comparing WT and respective bZIP mutants (for Fig. 1b, d, e, k, **l**, **n** see **Supplementary Table 4**), *** P<0.001.

Similar alterations in apical and lateral organ growth were observed during the reproductive phase. While bolting time of the primary inflorescence of the single and double mutants resembled that of the WT (Supplementary Fig. 1b), the triple and quintuple bZIP mutants exhibited a moderate or substantial delay, respectively (Fig. 1f-g). In addition, seed (Fig. 1h) and silique (Fig. 1i-j) growth at the primary inflorescence was significantly impaired in the mutants, whereas the latter could be rescued by removing all resource-consuming axillary and cauline branches that were about to grow out (Supplementary Fig. 1c). This indicated that silique growth at the primary, apical inflorescence was not generally defective in the mutants, but was likely short in resource supply.

In contrast to the impaired apical shoot growth responses, growth of lateral shoot organs was enhanced. The *bzip11* mutant for instance, showed a mild and transient increase of axillary branching, while the other single and double mutants showed no obvious growth phenotype (Supplementary Fig. 1d–h). In contrast, the triple (Fig. 1k) and quintuple (Fig. 1 j,l) mutants exhibited an early and strong enhancement of shoot branching.

Based on the *bzip11* branching phenotype and a strong *bZIP11* induction in leaf petioles and axillary buds upon establishment of the primary inflorescence (Supplementary Fig. 1i–j), we used this TF as a prototypical example for complementary gain-of-function approaches. To avoid artefacts caused by ectopic misexpression, we employed a dexamethasone (DEX)-inducible expression system, which allowed a rapid and tissue specific translocation of bZIP11 into the nucleus^30^. Compared to solvent (mock) control, repeated DEX application to the rosette core of *Pro35S*::*bZIP11*-*GR* plants severely suppressed axillary branching (Fig. 1m-n), while growth of the primary inflorescence remained unaffected (Supplementary Fig. 1k). In conclusion, the comprehensive phenotypic characterisation of the S_1_-bZIP loss– and gain-of-function lines clearly demonstrates a key function of S_1_-bZIPs in regulating apical and lateral shoot and root growth. Furthermore, a striking correlation between the order of S_1_ mutations and the reduction in apical dominance suggests a redundant function of these transcription factors (Fig. 1o).

### S_1_-bZIP TFs control leaf sugar export to the apoplast for long-distance transport to distal sinks

The promotion of lateral shoot and root growth at the expense of the respective apical organs and the rescue of impaired silique growth by removal of lateral sinks (branches) pointed towards a shifted nutrient allocation between apical and lateral organs in the S_1_-bZIP mutants. To test this hypothesis, we examined WT and bZIP mutants for their sugar content in primary roots and in apical and lateral shoot organs.

Concerning roots, we assessed C supply by determining the concentration of sucrose, the major sugar transport form, in primary roots of WT, bZIP triple and quintuple mutants. Strikingly, we found a roughly 50% reduction in root sucrose levels in the mutants (Fig. 2a), supporting the notion that S_1_-bZIPs are critical for efficient sugar channelling to the roots. To test whether the reduced sucrose supply caused the primary root growth repression, we performed sugar feeding experiments. Remarkably, cultivation of the triple and quintuple bZIP mutants on half-strength MS media supplemented with 30 or 60 mM sucrose, largely restored primary root growth to WT levels (Fig 2b-c, Supplementary Fig 2a-c). Notably, external sucrose supply also restored LR formation in a sucrose concentration-dependent manner (Supplementary Fig. 2d–l).

**Fig. 2.**
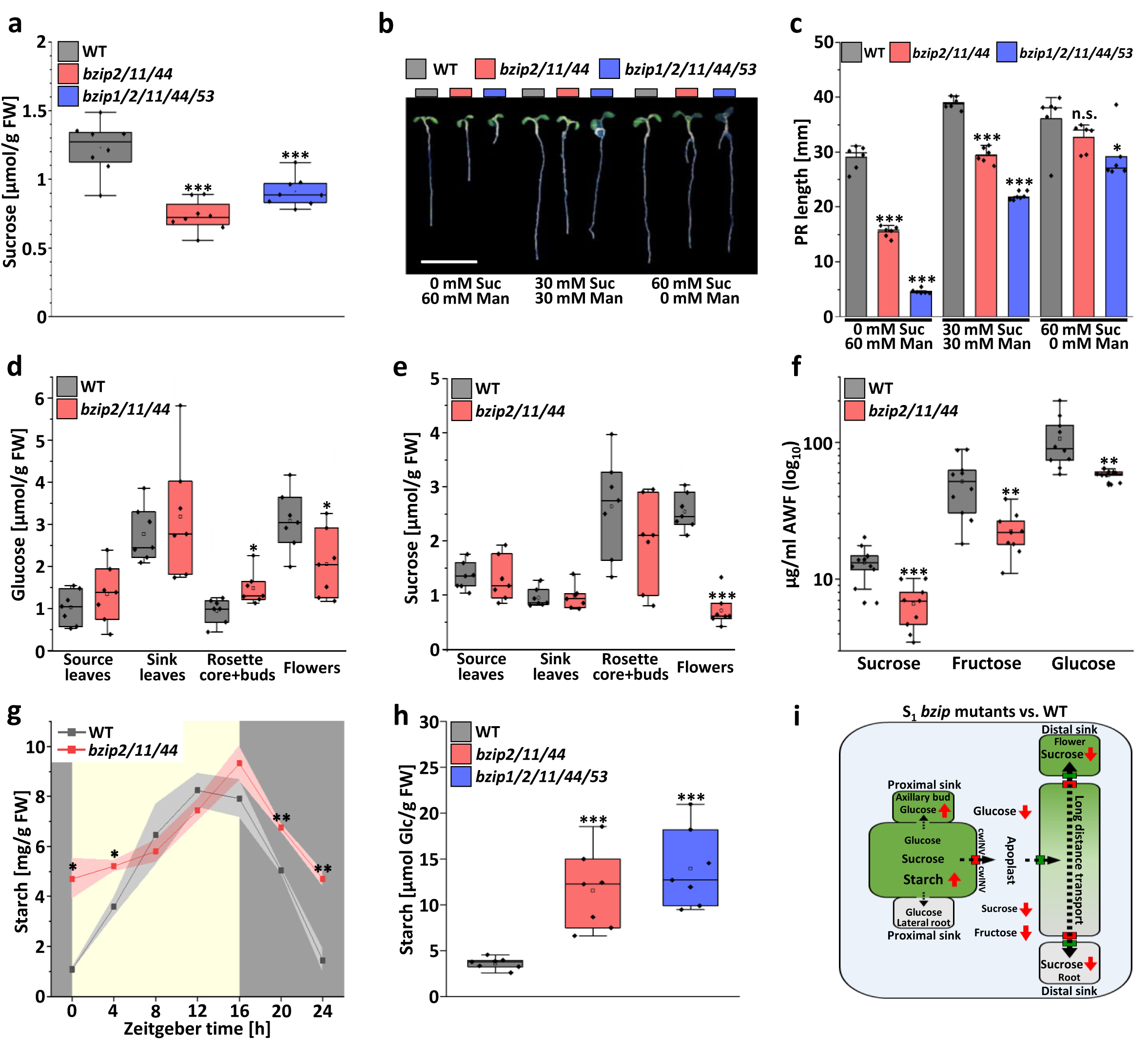
I S_1_ bZIP TFs are crucial for leaf sugar export to the apoplast for long distant transport to distal sinks. **a**, Primary roots of S_1_ bZIP mutants are sucrose depleted. Given is the mean root sucrose content (n=8 individual pools of roots per genotype) of 12-day-old WT (black box), bZIP triple (red box) and quintuple (blue box) mutants. **b-c**, Sucrose feeding can largely revert impaired root growth of S_1_ bZIP triple (red box) and quintuple (blue box) mutants back to WT (black box) level. Plants were grown for 10 days on ½ strength MS medium supplemented with 30 or 60 mM of sucrose (Suc) and/or mannitol (Man) as osmotic control. **b**, Representative picture of 7-day-old WT and bZIP mutant plants grown under control conditions or sucrose supplementation. Scale bar: 1 cm. **c**, Given is the mean primary root length (bar box plot, n=6, each replicate consisting of 10 plants per genotype) of WT (black box), bZIP triple (red box) and quintuple (blue box) mutants in presence or absence of sucrose. **d-e**, Distal sinks of S_1_ bZIP mutants are sugar depleted, while sugar supply of proximal sinks is not affected. **d**, Glucose and (**e**) sucrose content (n=7) was determined by UPLC-MS/MS in 8-week-old WT (black box) and bZIP triple mutants (red box). Stated tissues were harvested when primary inflorescence reached ∼10 cm. **f**, Leaf sugar export to the apoplast is strongly impaired in S_1_ bZIP mutants. Sugar (sucrose, fructose and glucose) content in apoplastic wash fluid (AWF) of WT (black box) and bZIP triple mutants (red box) was determined by UPLC-MS/MS. Whole rosettes of 6– week-old plants were vacuum-infiltrated with water and AWF was collected by centrifugation. Given are mean values (log_10_, n=9-10 per genotype) derived from AWF of two plants per replicate. **g-h**, Leaves of S_1_ bZIP mutants accumulate high levels of starch. **g**, Mean leaf starch content (± SEM, n=3 per genotype) throughout the day/night cycle (light/dark shaded area) of WT (black line) and bZIP triple mutant (red line). **h**, Mean starch content (n=7, each replicate consisting of 2 plant rosettes) in rosettes leaves of 5-week-old bolted WT (black box), bZIP triple (red box) and quintuple (blue box) mutants at ZT0. Starch was determined enzymatically. **i**, Schematic picture of sugar levels in distinct sink or source tissues of S_1_ bZIP mutants relative to WT (red arrows). Sugar transport routes from source to sink tissues (black arrows) as well as localisation of sugar export (red boxes) or import (green boxes) facilitators and cell wall invertases (cwINV) are given. Plants were either grown on soil (**Fig. 2d-h**) or ½-strength MS medium (**Fig. 2a-c**) under long day conditions at 100 µmol photons m^-2^ s^-1^. Statistically significant differences were determined by Student’s *t*-test comparing WT and respective bZIP mutants, n.s. not significant, *P < 0.05, **P < 0.01, ***P<0.001.

To define the sugar content in apical and lateral shoot organs (sinks) and shoot source tissues, we determined the levels of the two most abundant sugars – glucose (Fig. 2d) and sucrose (Fig. 2e) in mature leaves (source) as well as young leaves, rosette cores enriched in axillary buds, and flowers (sinks) at an early reproductive stage, prior to branching. Due to the extremely early branching phenotype of the quintuple mutant, we focused our analysis on WT and triple mutants. While we did not observe any difference in glucose or sucrose levels in leaves of either genotype (Fig. 2d-e), we detected a strong decrease in distal sink tissues (flowers) of the triple mutant (Fig. 2d–e). In contrast, triple mutants showed a significant increase in glucose levels in proximal sinks (rosette core with axillary buds) compared to WT plants (Fig. 2d). Therefore, we hypothesised that leaf sugar export is impaired in the triple mutant, which would increase the symplastic sugar flux to proximal sinks but inhibit long-distance transport through the apoplast and vasculature to distal sinks. To further pursue this, we isolated leaf apoplastic wash fluid from whole rosette leaves and measured the concentration of sucrose and its degradation products glucose and fructose that are produced by apoplastic, cell wall-associated invertases (cwINV). Consistent with an impaired export from mesophyll cells, apoplastic sugars were severely reduced by about 50% in the mutant (Fig. 2f).

Impaired sugar export from leaf cells should ultimately favour starch biosynthesis to counteract the toxicity of high intracellular hexose concentrations. Accordingly, we found significantly elevated starch levels in the triple mutant (Fig. 2g), especially during the night and the beginning of day. A slightly stronger effect was also observed in the quintuple mutant at dawn (Fig. 2h). From these results, we conclude that group S_1_– bZIP TFs operate in leaves to support sugar export to the apoplast and, consequently, long-distance transport to distal sinks, such as the apical shoot and root meristems (Fig. 2i).

### Shoot and root glutamine supply is controlled by S_1_-bZIP TFs

Besides C availability, supply of organic N is essential for meristem activity. Under low– energy conditions, individual group S_1_-bZIPs were found to induce degradation of proline and the branched chain amino acids leucine, isoleucine, and valine to feed carbon skeletons into alternative respiratory pathways for ATP production^23^. To obtain a comprehensive picture of the organic N status during development of the higher-order bZIP mutants under non-starved conditions, we determined the levels of proteinogenic AAs in shoot and root tissues of WT, bZIP triple and quintuple mutants in the middle of the day at an early and rather late vegetative stage. In contrast to what was found in response to energy depletion, we did not observe a considerable change in proline and branched chain AA levels throughout tissue development of the genotypes (Fig. 3a-c, Supplementary Fig. 3). However, a striking increase in the levels of the most abundant AAs Gln and asparagine (Asn), which serve as major organic N transport forms during the day or night, respectively^32^, was detected in seedling roots of the triple (Fig. 3a) and, to an even greater extent, in the quintuple mutant (Supplementary Fig. 3) compared to WT. Although, the effect on transport AAs was not found in the shoot tissue of the triple mutant at the seedling stage (Fig. 3b) a clear and specific increase in Gln levels was observed just before bolting (Fig. 3c). Therefore, we propose that group S_1_-bZIP TFs redundantly act in roots to systemically control homeostasis of the central organic N transport form Gln during plant development.

**Fig. 3.**
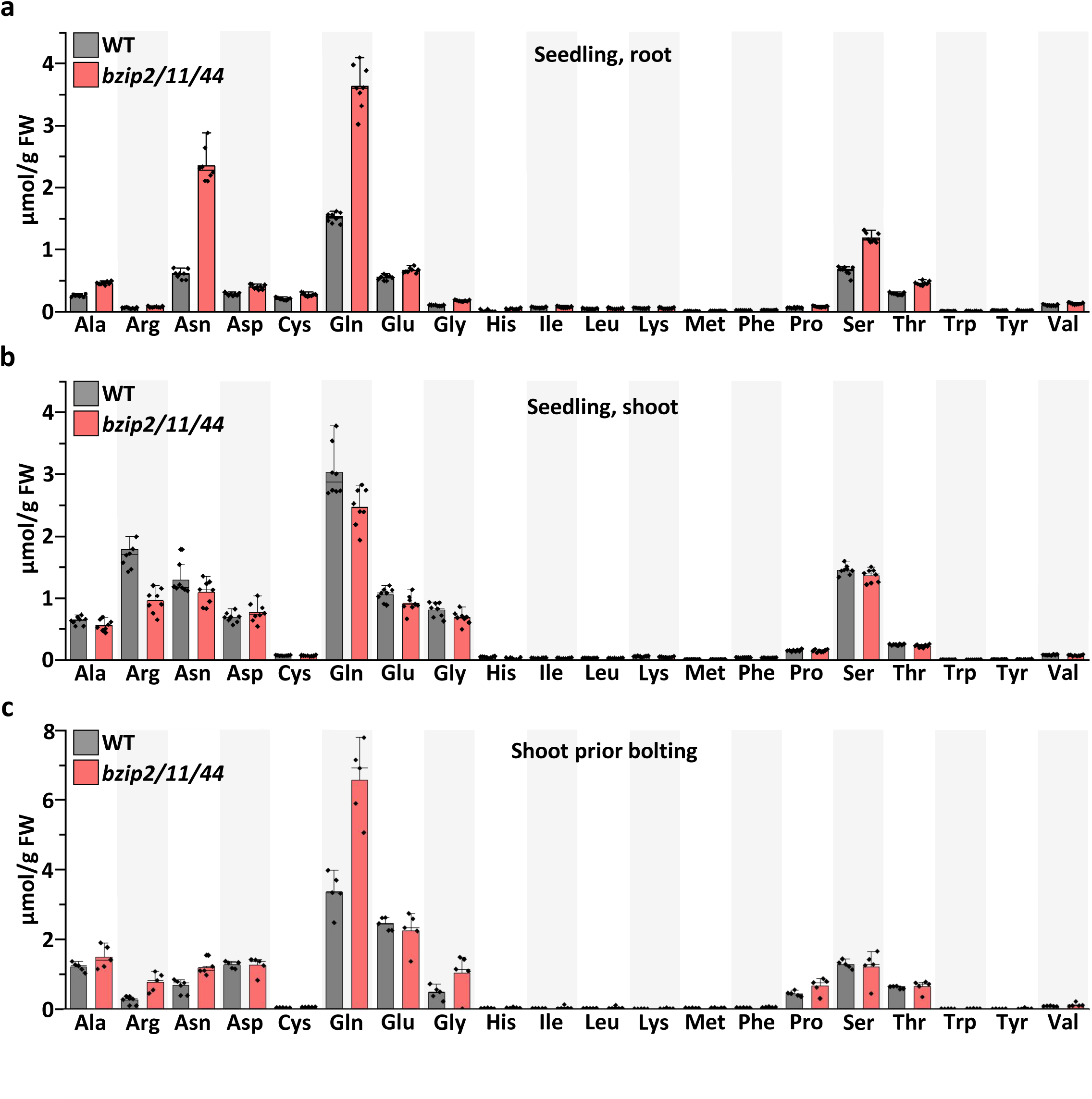
I S_1_ bZIP mutants show a marked increase in glutamine levels in roots and shoots prior bolting. **a**, Given is the mean amino acid content (bar box plots, n=8) determined by UPLC-MS/MS from roots of 2-week-old WT and bZIP triple mutant seedlings. Root material of several plants was harvested at ZT8 and pooled. **b**, Mean amino acid content (bar box plots, n=8) in shoots of 2-week-old WT (black box) and bZIP triple mutant (red box) seedlings. Shoot material of several plants was harvested at ZT8 and pooled. **c**, Mean amino acid content (bar box plots, n=5, each replicate consisting of 2 plants) from rosettes of 4-week-old WT or bZIP triple mutant plants harvested at ZT8. Plants were grown on (**a**, **b**) ½ strength MS medium or (**c**) soil under long-day conditions at 100 µmol photons m^-2^ s^-1^. Statistically significant differences were determined by Student’s *t*-test comparing individual amino acid content of WT and respective bZIP mutants (see **Supplementary Table 4).**

### S_1_-bZIP TFs control tissue-specific expression of defined clade 3 *SWEET* sugar transporters and of the glutaminase *GAT1_2.1*

To gain insight into the mechanisms by which S_1_-bZIPs control local C/N supply, we performed RNA-sequencing (RNAseq) to identify putative S_1_ targets in source and sink tissues. To visualise the complex dataset (Supplementary Table 2), we generated Venn diagrams showing the significantly (p<0.05) up– (log_2_FC g 1) or down-regulated (log_2_FC f –1) genes of the triple (Fig.4 a-b) or quintuple (Fig.4 c-d) mutant relative to WT plants. Consistent with previous work revealing individual S_1_-bZIPs as transcriptional activators recruiting the histone acetylation machinery^33^, we generally found more down-than upregulated genes in the triple and quintuple mutants (Fig. 4a–d). In addition, we observed a high degree of tissue specificity, suggesting sink/source-specific regulation of S_1_ factors. Importantly, several sugar transporters of the SWEET family, known to facilitate leaf sugar export^34^, were found among the differentially expressed genes (DEGs) (Fig. 4e-f). Of these, predominantly clade 3 *SWEETs* (*SWEET9-15*)^11^ were affected, with *SWEET10*, *-13*, and *-14* strongly down-regulated in sinks and *SWEET15* down-regulated in both source and sink tissues (Fig 4e-g). To address whether S_1_-bZIPs directly control their expression, we analysed a publicly-available DNA affinity purification sequencing (DAPseq) database^35^ to examine direct, *in vitro* promoter binding. In addition, we performed an assay for transposase-accessible chromatin by sequencing (ATACseq) from protoplasts of transgenic plants that were transfected with a DEX-inducible bZIP11 construct and were treated with the translation inhibitor cycloheximide to uncover direct, *in vivo* bZIP11-dependent chromatin opening of *SWEET* promoter regions. While binding of S_1_-bZIPs was observed at all tested *SWEET* promoters (Supplementary Fig. 4a–d), only *SWEET14* and especially *SWEET15* exhibited a strong bZIP11-dependent chromatin opening at the respective sites (Supplementary Fig. 4e–h). Applying protoplast transactivation assays, we validated the transactivation properties of S_1_-bZIPs on the highly responsive *SWEET15* promoter (Fig. 4h) and pinpointed two cognate bZIP *cis*-elements (Gbox-related elements, GREs) as critical for promoter transactivation (Fig. 4i).

**Fig. 4.**
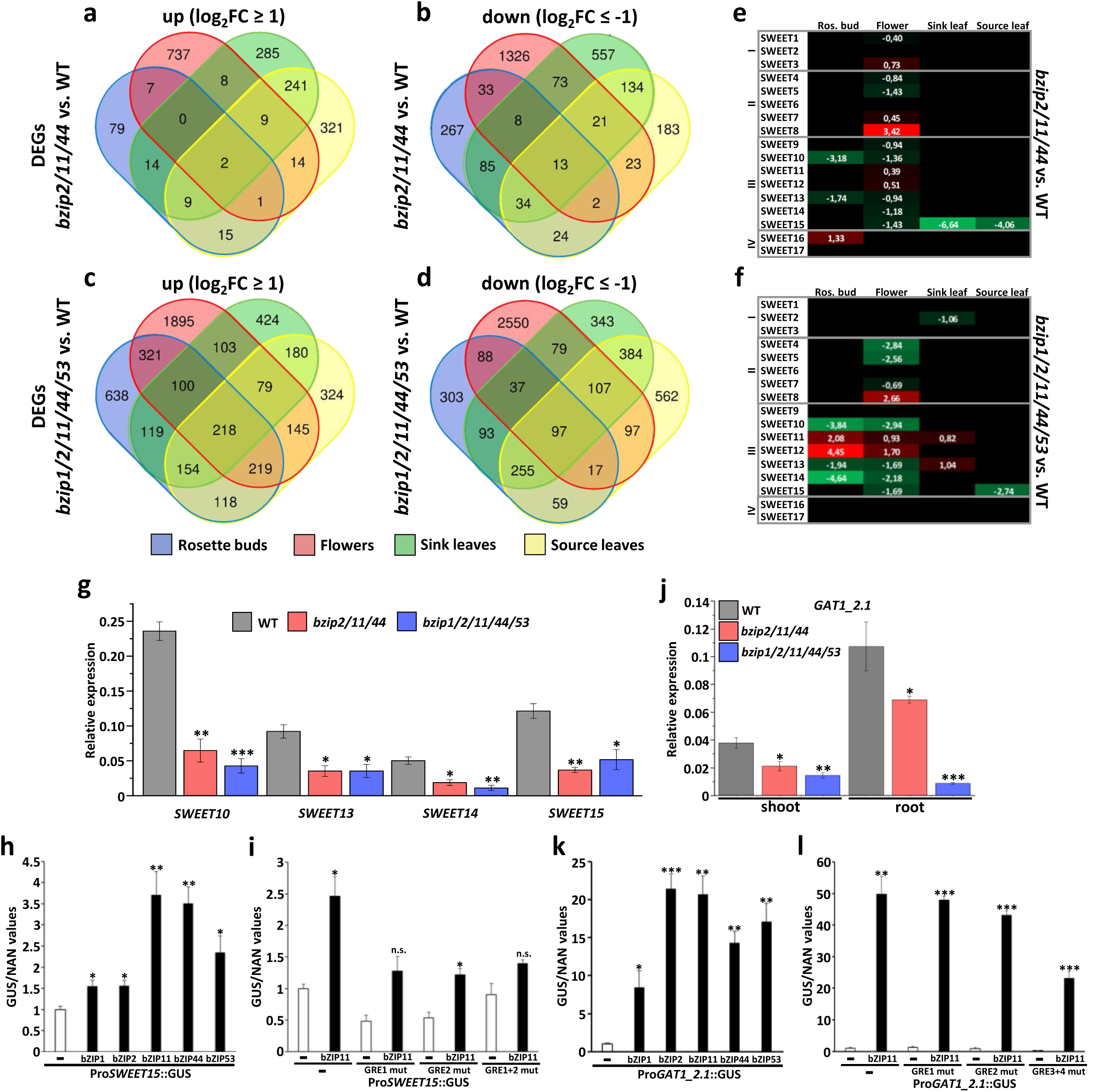
I S_1_ bZIP TFs control expression of *SWEET15* and *GAT1_2.1* in distinct plant tissues. **a-d**, Differentially expressed genes (DEGs) between WT and S_1_ bZIP mutants in distinct C source and sink tissues. Global transcript abundance in specific tissues (rosette buds, blue; open flower, red; young leaves, green; and old leaves, yellow) of WT, bZIP triple and quintuple mutant plants at pre-branching state was determined by RNA sequencing (n=3 per genotype and tissue). Venn diagrams present significantly (p < 0.05) up (log_2_FC g 1) (**a**,**c**) or down (log_2_FC f –1) (**b**,**d**) regulated genes comparing WT, bZIP triple or quintuple mutant plants, respectively. Complete list of DEGS can be found in Table S2. **e-f**, Overview of tissue specific expression of *SWEET* family members (grouped in clades I to IV) in (**e**) bZIP triple or (**f**) bZIP quintuple mutants relative to WT. Given are log2-fold expression changes (induction given in shades of red, repression in shades of green). **g**, Validation of *SWEET1O*, *-13*, *-14* and *-15* expression by quantitative Realtime PCR (q-RT-PCR) in flowers of WT (black bar), bZIP triple (red bar) or quintuple (blue bar) mutants prior branching. Presented are mean expression values (± SEM, n=3) relative to UBIQUITIN5 expression. **h-i**, Protoplast transactivation assays (PTAs) using a *SWEET15* promoter driven GUS reporter (Pro*SWEET15*:GUS) and Pro35S driven S_1_ bZIP effector constructs. Pro35S driven NAN expression was used for normalisation. **h**, *SWEET15* promoter activity in absence (**-**) or presence of bZIP effector constructs (bZIP1,-2,-11,– 44,-53) given as mean GUS/NAN values (± SEM, n=3) relative to promoter background levels. **i**, Promoter activity of the original and GRE mutated *SWEET15* promoter in absence (**-**) or presence of bZIP11 effector construct given as mean GUS/NAN values (± SEM, n=2) relative to promoter background levels. **j**, Expression analysis of *GAT1_2.1* in shoots and roots of 2-week-old WT (black bar), bZIP triple (red bar) or quintuple (blue bar) mutants by q-RT-PCR. Presented are mean expression values (± SEM, n=3) relative to UBIQUITIN5 expression. **k-l**, PTAs using a *GAT1_2.1* promoter driven GUS reporter (Pro*GAT1_2.1*:GUS) and Pro35S driven S_1_ bZIP effector constructs. Pro35S driven NAN expression was used for normalisation. **k**, *GAT1_2.1* promoter activity in absence (**-**) or presence of bZIP effector constructs (bZIP1,-2,-11,-44,-53) given as mean GUS/NAN values (± SEM, n=3) relative to promoter background levels. **l**, Promoter activity of the original and GRE mutated *GAT1_2.1* promoter in absence (**-**) or presence of bZIP11 effector construct given as mean GUS/NAN values (± SEM, n=2) relative to promoter background levels. Plants were either grown on soil (**a-I**,**k-l**) or ½-strength MS medium (**j**) under long day conditions at 100 µmol photons m^-2^ s^-1^. Statistically significant differences were determined by Student’s *t*-test comparing WT and respective bZIP mutants, n.s. not significant, *P < 0.05, **P < 0.01, ***P<0.001.

To identify targets that are causative for the increased root Gln levels in S_1_-bZIP mutants, we screened a previously published transcriptomics dataset of *bZIP11* overexpression lines^30^. Among the DEGs, the glutaminase *GAT1_2.1* emerged as the most promising candidate as it 1.) was strongly induced by bZIP11^30^, 2.) was predominantly expressed in roots^15^, and 3.) showed a broad binding of S_1-_bZIPs in a GRE-rich promoter region (Supplementary Fig. 4i), which also exhibited a significant bZIP11-dependent increase in chromatin accessibility (Supplementary Fig. 4j). Consistent with these results, we found a strong downregulation of root *GAT1_2.1* expression in the bZIP triple and, to an even greater extent, in the quintuple mutant (Fig. 4j). We also observed a strong and redundant activation of the *GAT1_2.1* promoter by S_1_-bZIPs (Fig. 4k), which was at least partially accomplished by two adjacent GREs located within the proposed bZIP11 binding site (Fig. 4l).

### Mutants of S_1_-bZIP targets exhibit a shoot branching phenotype

TFs generally control a plethora of target genes. To test whether the addressed clade 3 *SWEETs* and the *GAT1_2.1* glutaminase contribute significantly to the S_1_-bZIP-driven repression of lateral organ growth, we monitored axillary branching of loss-of– function mutants of the respective target genes. While we observed no increase in shoot branching between WT plants and an unrelated *sweet8* mutant (Supplementary Fig. 5a), *sweet15* mutants exhibited an early branching phenotype (Fig. 5a-b). Single *sweet10*, *sweet13*, *sweet14* as well as double *sweet13/-14* mutants showed a moderate but consistent increase in the number of axillary branches (Supplementary Fig. 5b–d). Similarly, *gat1_2.1* mutants showed a significant increase in shoot branching (Fig. 5c-d).

**Fig. 5.**
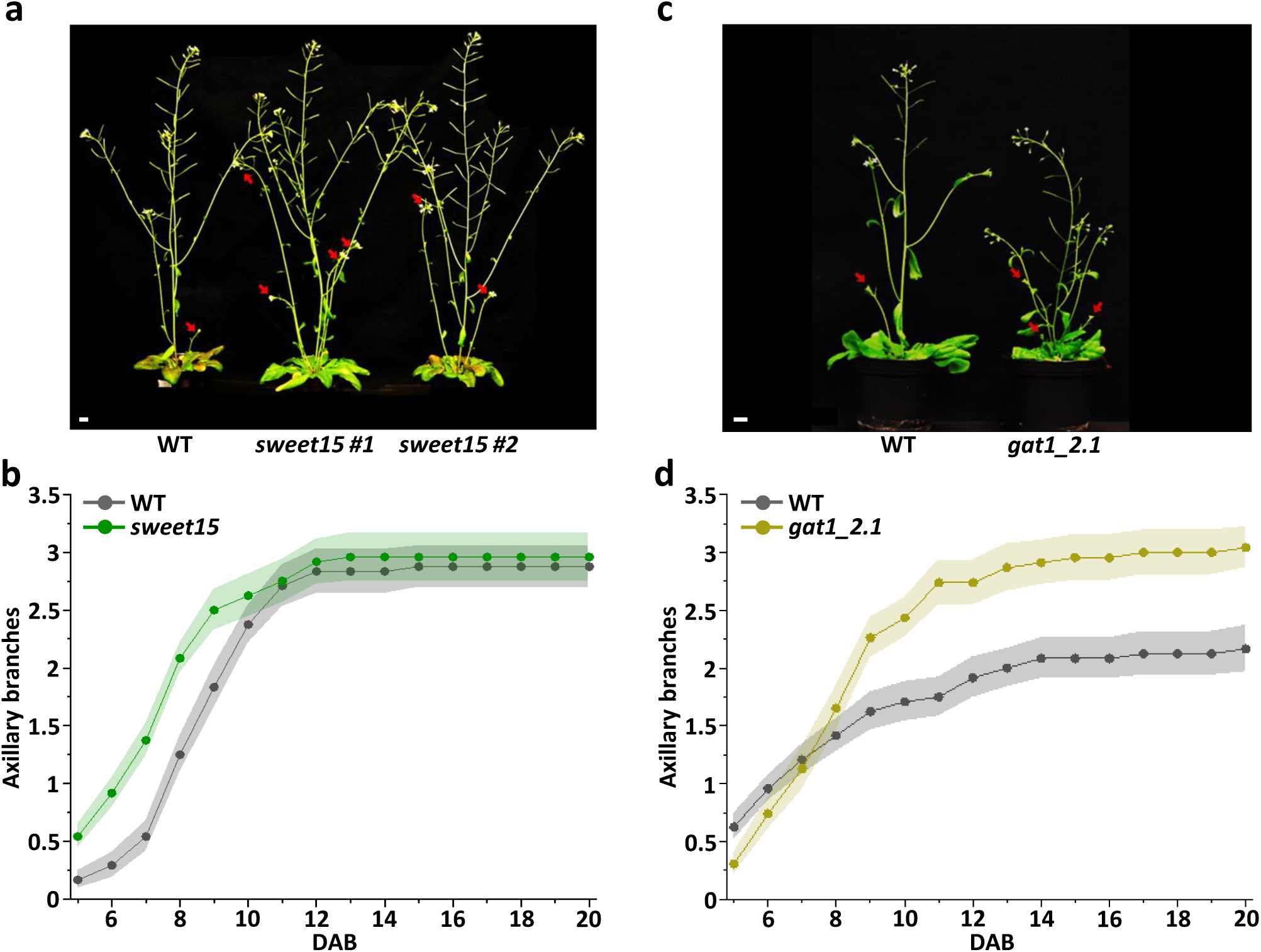
I Mutants of *SWEET15* and *GAT1_2.1* reveal a high branching phenotype. **a-d**, Axillary branching of *SWEET15* and *GAT1_2.1* mutants. **a**, Representative picture of 6-week-old WT and two independent *sweet15 plants.* Red arrows mark outgrown axillary branches. Scale bar: 1 cm. **b**, Mean axillary branching (± SEM, n=24 per genotype) of WT (black line) or *sweet15* (green line) plants. **c**, Representative picture of 5-week-old WT and *gat1_2.1 plants.* Red arrows mark outgrown (g1cm) axillary branches. Scale bar: 1 cm. **d**, Mean axillary branching (± SEM, n=24 per genotype) of WT (black line) or *gat1_2.1* (yellow line) plants. Plants were grown on soil under long day conditions at 100 µmol photons m^-2^ s^-1^. Outgrown axillary branches (g1cm) were counted starting 5 days after bolting (DAB). Statistically significant differences were determined by Student’s *t*-test comparing WT and respective mutants (see **Supplementary Table 4)**.

## Discussion

Plants display an enormous developmental plasticity in shaping shoot and root architecture. Although most plants show a pronounced apical growth dominance, lateral organ formation is inducible and highly sensitive to the plant’s sugar status^6, 9^. Recent work demonstrated that rapid sugar reallocation to dormant axillary branches upon decapitation acts as an initial trigger for bud outgrowth^5^ and highlight the impact of sugar signalling in shoot branching^36–39^. This suggests that plants possess a sophisticated mechanism for sensing nutrient availability and allocating limited resources between apical and lateral meristems to control their growth.

Initially characterised as central players in the plant’s energy management system^25^, we unravelled in this study a novel, developmental function of the subgroup S_1_-bZIP TFs (bZIP1, –2, –11, –44 and –53) in tuning apical growth dominance. Known to be expressed under energy-depleted conditions^40^ and in strong sink tissues (such as flowers^29^, seeds^41^ and roots^26^), we also observed bZIP11 expression in petioles of source leaves upon establishment of the primary inflorescence sink (Supplementary Fig. 1i–j). This suggests that S_1_-bZIP expression is controlled by either tissue-specific or developmental signals, if not both. This hypothesis is supported by the RNAseq datasets derived from source and sink tissues of the S_1_-bZIP mutants (Fig. 4 a-d), which show a high degree of tissue specificity in bZIP-mediated expression. Unexpectedly, the typical starvation responsive targets of S_1_-bZIPs (e.g. *ETFQO*, *MCCA* etc.)^23^ are not differentially expressed in the analysed tissues, suggesting a clear difference in bZIP action in starvation-related or developmental processes.

Consistent with the marked decrease in sugar supply to the leaf apoplast and apical sink tissues (Fig. 2a, d-f), and sugar increase in leaf cells and proximal sinks such as the axillary branches (Fig. 2d, g-h), we observed a shift from apical growth promotion to lateral organ formation in the S_1_-bZIP mutants (Fig. 1a-l). This pointed towards a coordinated change in sugar partitioning to the apical sinks. In line with this, sugar supplementation of the C-depleted roots or reduction in C-demand of the shoot by excision of cauline branches rescued apical growth phenotypes of the bZIP mutants such as primary root growth or silique expansion on the primary shoot inflorescence (Fig. 2b-c, Supplementary Fig. 1c). Examination of protoplast transactivation assays (Fig. 4h-i), RNAseq (Fig. 4e-f), ATACseq and DAPseq^35^ datasets (Supplementary Fig. 4a–h) identified specific clade 3 SWEET sugar transporters as critical targets in this regard. Consistently, the single *sweet10*, *sweet13*, *sweet14*, *sweet15* and the double *sweet13/-14* mutants showed reduced apical dominance (Fig. 5a-d, Supplemental Fig. 5a-d), and the latter also had shortened siliques and increased lateral root formation^42^, partially resembling S_1_-bZIP mutant phenotypes (Fig. 1i-l). Higher-order mutants of clade 3 *SWEETs* could further assist in unravelling their contribution to S_1_-bZIP driven apical dominance. However, due to prevalent SWEET heterodimerisation^43^, these mutants would likely exhibit highly pleiotropic effects. While the discussed *SWEETs* are coordinately downregulated in sink tissues of the bZIP mutants, *SWEET15* expression was also affected in leaves, likely leading to impairment of both sugar export from the source and import at apical sink tissues (Fig. 4e-f) and consequently reduced apical growth. Importantly, both *SWEET* expression and sugar translocation to apical meristems are equally impaired in bZIP triple and quintuple mutants, indicating that bZIP2, –11 and –44 are particularly crucial in this respect (Fig. 2a,h; 4g).

In addition to their function in C partitioning, we also uncovered the impact of S_1_-bZIPs on Gln homeostasis. Representing an important reservoir for organic N, Gln is essential for organ growth and development. In this respect, we observed a clear correlation between the order of S_1_-bZIP mutations, the increase in root Gln levels (Fig. 3b, Supplementary Fig. 3) and the expression of lateral organ growth (Fig. 1e, Supplementary Fig. d,g,j). It is worth noting that Gln levels in the mutants were only elevated in roots and not in shoot tissues at the early vegetative stage (Fig. 3a-b), while a strong increase was observed in the shoot at the onset of the reproductive phase. This suggests that either root Gln is efficiently transported to the shoot upon bolting and/or that again bZIP action is subject to tissue– and development-specific regulation. Making use of protoplast transactivation assays (Fig. 4k,l), transcript analysis by qRT-PCR (Fig. 4j), ATACseq and DAPseq^35^ data (Supplementary Fig. 4i,j), we identified the recently described glutaminase *GAT1_2.1*^14^ as a direct target of S_1_-bZIPs. Consistently, largely overlapping expression domains of *GAT1_2.1* and *S_1_-bZIPs* were revealed in sink tissues, such as flowers, siliques, axillary branches, and roots^15, 29^. In line with this, all S_1_ factors appear to contribute to the full expression of *GAT1_2.1*, which is most severely downregulated in the quintuple S_1_ mutant (Fig. 4j). Complementary to the loss-of-function approach, inducible bZIP11 overexpressors showed a strong and rapid activation of *GAT1_2.1* transcription, which was accompanied by a marked decrease in plant Gln levels^30^. Furthermore, DEX-application to the rosette core of this line, revealed a strong bZIP-dependent repression of axillary branching (Fig. 1m,n). Conclusively, phenotypic analysis of a *gat1_2.1* mutant validated its impact on shoot branching (Fig. 5c-d)^15^.

In summary, we propose that S_1_-bZIPs have a dual function in establishing apical dominance, by controlling C and N allocation to apical and lateral organs (Fig. 6). Based on our results we propose that this is accomplished by S_1_-bZIP TFs that control *SWEET* expression in source (leaf petioles) and distal sink tissues (shoot and root apical meristems) to generate a constant sugar flow to the growing apical organs, while keeping the lateral primordia in a sugar depleted state. The central impact of S_1_-bZIPs on root glutaminase abundance and in consequence global organic N supply further assists in tuning organ development.

**Fig. 6.**
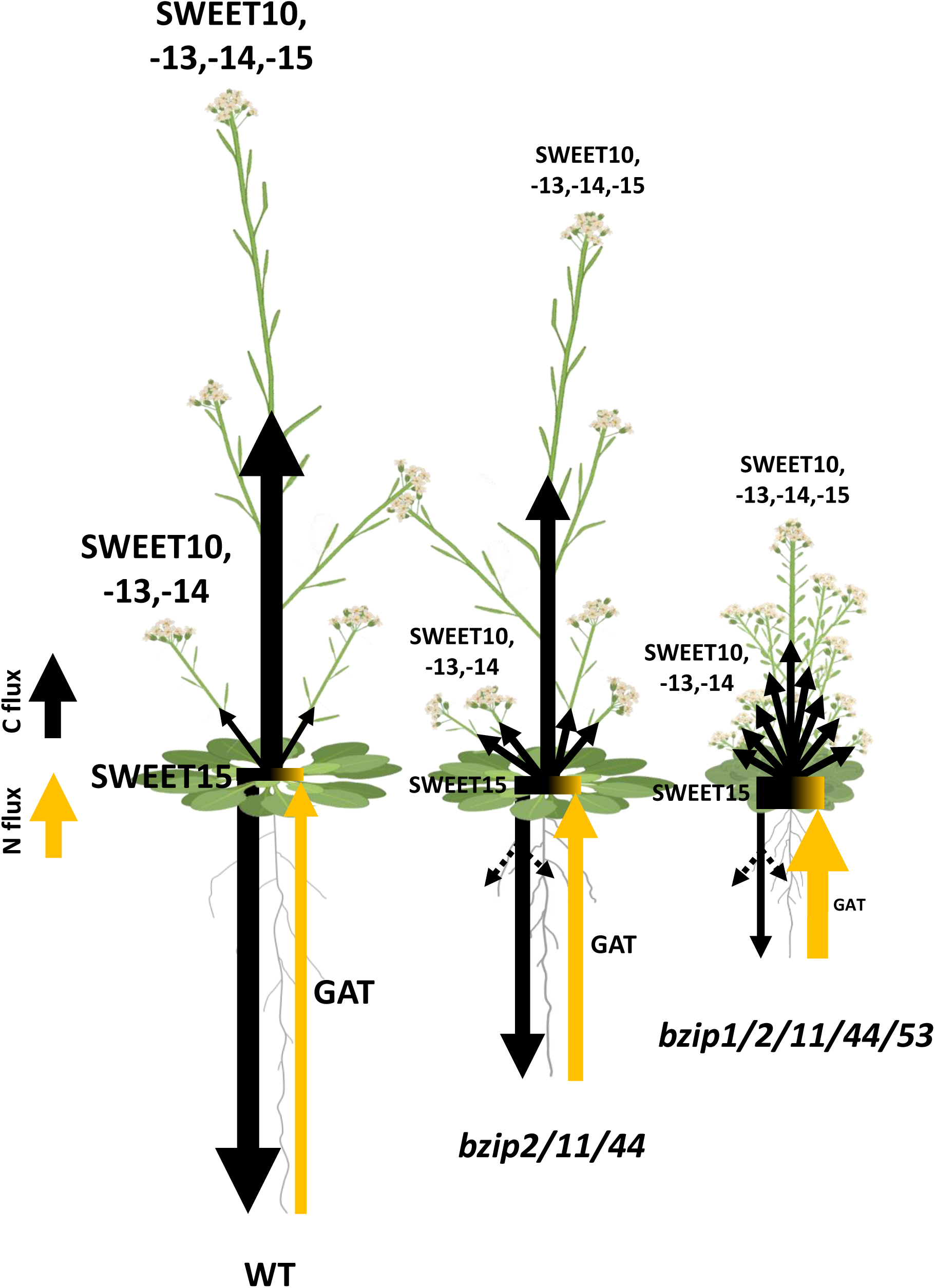
I Model of S_1_ bZIP action in steering C and N partitioning to control shoot and root architecture. The model plant Arabidopsis as well as many agronomically important plants species display a pronounced apical growth dominance, while lateral organs remain quiescent until the sink demand of the primary meristem declines. To enable such a differential growth control of apical shoot / root meristems and the lateral shoot (axillary branch) / root (lateral root) primordia, plants require a mechanism to direct fluxes of C (sugars, black arrows) and N resources (amino acids, yellow arrow) from places of production (source tissues) to the specific resource-consuming organs that are destined to grow (sink tissues). This is accomplished by coordinated sugar and AA partitioning. While sucrose efflux transporter family proteins (SWEETs) facilitate passive diffusion of photosynthesis-derived sugars from rosette leaves to the leaf apoplast, sugar transporters of the SUC/SUTs family actively load apoplastic sugars against the gradient into the phloem. Predominantly at strong apical sinks, like the flowering shoot apex or the growing primary root meristem, SWEETs again assist in unloading of phloem sugars, thereby creating a constant sugar gradient towards the apical meristems. In reverse orientation, glutamine, which serves as a transport form of root-fixed inorganic N is loaded into the root xylem to be transported to the actively growing shoot and root sinks. As S_1_ bZIPs directly and redundantly control expression of central clade 3 *SWEETs* at source (leaf) and sink (flower) tissues and of root GLUTAMINE AMIDOTRANSFERASE 1_2.1 (*GAT1_2.1*) that is known to initialize glutamine degradation, they directly control C canalization to the primary meristems and balance systemic N homeostasis. Consistently, higher-order mutants of S_1_ bZIPs are disturbed in long distance sugar transport and display enhanced leaf starch accumulation and diffusion of hexoses, likely via symplastic routes, to proximal sinks, such as axillary branches and lateral roots. An additional deregulation of root glutamine homeostasis in the mutants, results in a marked increase in shoot and root N supply providing all resources required for lateral meristem outgrowth. In conclusion S_1_ bZIPs play a pivotal role in managing C/N partitioning to promote apical but restrict lateral meristem activity.

As S_1_-bZIPs are highly conserved in agronomically relevant plants, manipulation of this system would help to create plants with increased or reduced branch number, which would be valuable for fruit, flower and timber industries. In addition, local misexpression could be used to direct sugar fluxes to specific tissues. Consistently, recent reports have shown that fruit-specific misexpression of S_1_-bZIP orthologs in strawberry or tomato results in a strong increase in fruit sugar content^44–46^. Thus, our studies broaden the view on the developmental impact of the stress-related S_1_-bZIP TFs to control energy-intensive plant architecture by determining the C/N status of meristems, and open a gateway for biotechnological progress.

## Methods/ Online methods

### Transgenic lines, plant cultivation and phenotypic characterisation

In all assays WT *Arabidopsis thaliana* accession Columbia (Col-0) was used as control, and all transgenic lines in this study are in a Col-0 background. The s*weet13* (SALK_087791, psi00187), *sweet14* (SALK_010224, psi00188) and *sweet13 x sweet14* (psi00189) mutants were characterised previously^42^ and were obtained from RIKEN (https://www.riken.jp). The s*weet15* mutant line was characterised and kindly provided by the Coupland lab. The *gat1_2.1* (SALK_031983C, N665686) mutant was characterised previously^14^ and was obtained from NASC (https://arabidopsis.info/).

Pro*bZIP11*::*bZIP11*-GFP and Pro*bZIP11*::*bZIP11*-GUS plant lines were constructed by PCR-amplifying 2.3 kb of the *bZIP11* promoter and the joining *bZIP11* coding region without stop from genomic DNA using Gateway^®^-site flanking primers, given in Supplementary Table 3. PCR-products were initially integrated into the pDONR207 donor vector by Gateway^®^ BP-reaction (Thermo Scientific) and then shuttled to the binary plant expression vectors pGWB550 (c-term GFP)^47^ or pGWB533 (c-term GUS)^47^ using the Gateway^®^ LR clonase mix. Binary vectors were used for floral-dip transformation and resulting homozygous lines for experimental approaches.

The *sweetB*, *sweet10* and all single and higher-order mutants of S_1_-bZIP TFs were generated in this work using an egg-cell specific CRISPR/Cas9 genome editing system^31^. Target specific guide sequences positioned close to the translational start codon, ATG, were obtained from ChopChop^48^ and are given in Supplementary Table 3. Transgenic lines were selected by hygromycin treatment. Homozygous mutants, exhibiting frame-shifts and consequently premature stop codons in *SWEET* and *bZIP* genes were identified by sequencing PCR products of the putatively mutated regions. Genotyping primers used for this step are listed in Supplementary Table 3. An overview of mutations in the respective mutants is given in Supplementary Table 1.

Prior to sowing on growth substrate, seeds were surface sterilised with chlorine gas and stratified at 4°C for 3 days to synchronise germination. If not otherwise specified, plants were grown on soil under a long-day regime (16h light / 8 h darkness) at 21°C, an irradiance of 100 µmol photons m^-2^ s^-1^, and a relative humidity of 60%. For all assays addressing seedling shoot and root metabolism and root growth responses plants were cultivated aseptically on 1% (w/v) agar plates with ½-strength Murashige and Skoog medium.

To phenotypically characterise mutants, plants were grown side-by-side with WT controls under the specified growth conditions. Vegetative growth was assessed on a regular basis (every 2-3 days) by measuring leaf area using the BlattFlaeche tool (Datinf GmbH, Germany). Bolting time of the primary inflorescence (inflorescence g1cm) and axillary branching (branches g1cm) were scored daily. Average silique length of siliques 6-15 from the inflorescence base was determined at the end of the reproductive phase when siliques started to dry out. Seed weight was assessed by weighing individual batches of a defined number of seeds. From this, the average weight of a single seed was calculated. Root parameters such as PR growth, number of LRs and LR density given as <emerged LR per cm PR over time= or <LR count over PR growth= was determined by daily measuring of PR length and counting macroscopically visible LRs using the Fiji software^49^.

### Western Blot analysis

To determine bZIP11 protein amount in the axillary bud enriched rosette tissue at bolting or anthesis, three individual rosette cores of a *ProbZIP11*::*bZIP11*-*GFP* line and WT controls were harvested at the respective developmental stage and pooled to create a biological replicate. From three biological replicates of the transgenic line and two of WT plants, proteins were extracted and used for SDS-PAGE and subsequent western blotting. bZIP11-GFP was detected using a rat monoclonal α-GFP antibody (3H9; Chromotek; https://www.ptglab.com) at 1:2000 dilution. Histone 3 detection using a rabbit polyclonal α-H3 antibody (AS10710; Agrisera; https://www.agrisera. com) at 1:5000 dilution served as loading control.

### Histological GU5 staining

To visualise bZIP11-protein in rosette cores, we used *ProbZIP11*::*bZIP11*-*GUS* lines and performed histological GUS staining according to previously published work^50^ with minor modifications. In brief, plant material was fixed in 90% (v/v) acetone for 20 min at room temperature and afterwards washed with rinsing buffer (50 mM NaPO_4_, pH7.2; 0.5 mM K_3_Fe(CN)_6_, 0.5 mM K_4_Fe(CN)_6_). Subsequently tissue was transferred to staining solution (50 mM NaPO_4_, pH7.2; 0.5 mM K_3_Fe(CN)_6_, 0.5 mM K_4_Fe(CN)_6_, 5mM X-Gluc). To facilitate infiltration of buffer into the tissue, a vacuum was applied and released several times. Tissues were incubated in staining buffer over night at 37°C before they were treated with an ethanol series from 15% to 100% (v/v) to increase signal to noise ratio by removing chlorophyll.

### Sugar measurements

Unless stated otherwise, sugar (glucose, fructose, sucrose) measurements were performed as published previously^20^, with minor modifications. In brief, around 20 mg (fresh weight) of individual tissues (roots, leaves, rosette cores with branches or flowers) were harvested at ZT0 and frozen in liquid N_2_. Sugars from ground tissue were extracted with 300 µL 80% (v/v) ethanol, containing 2 µg [1,1-^2^H_2_]-trehalose and 8 µg [6,6-^2^H_2_]-glucose as internal standards. Samples were incubated at 80°C for 20 min and centrifuged for 10 min at 18,000 x *g*. After transferring the supernatant to a new reaction tube, the pellet was reextracted twice, first with 300 µL 50% (v/v) ethanol and, subsequently with 300 µL 80% (v/v) ethanol both at 80°C for 20 min. The extracts were pooled, and the solvent completely evaporated using a vacuum concentrator at 45°C. For MS (UPLC-MS/MS) analysis the obtained pellet was redissolved in 25 µL 50% (v/v) methanol.

Apoplastic sugars were extracted based on a previously published method^51^ with minor modifications. In brief, plants were grown on soil under long-day conditions at 100 µmol photons m^-2^ s^-1^ for six weeks. Rosettes of two individual plants per genotype were harvested and washed multiple times with ice-cold water. Afterwards rosettes of the same genotype were transferred to 60 ml plastic syringes. Syringes were filled with ice-cold water, which was carefully ejected until the 40 ml mark, thereby removing all air from the syringe. After the tip was sealed with parafilm, the plunger was slowly pulled back to the 60 ml mark to create a vacuum, which was then slowly released again. The air pulled from the leaves was ejected and the syringe sealed again. A slight positive pressure was applied to the plunger to improve water infiltration of leaves. If necessary, the process was repeated until the entire leaf area was infiltrated. Rosettes were then carefully dried using tissue paper and placed in a new 50 ml tube. Samples were centrifuged at 1500 x *g* and 4°C for 20 min. Rosettes were removed from the tubes and the remaining apoplastic wash fluid was collected. Of this 300 µl were lyophilized overnight. Samples were dissolved in 25 µl of 50% (v/v) acetonitrile containing 100 µg/ml [1,1-^2^H_2_]-trehalose and 400 µg/ml [6,6-^2^H_2_]-glucose as internal standards.

To determine sugar content 5 µL of the samples were analyzed using a Waters Acquity ultrahigh-performance liquid chromatograph coupled to a Waters Micromass Quattro Premier triple quadrupole mass spectrometer (Milford) with electrospray interface (ESI). Chromatographic separation was performed according to application note WA60126 with a modified flow rate of 0.2 mL/min. Sugars were detected in negative electrospray mode (ESI−) at 120°C source temperature and 3.25 kV capillary voltage. Nitrogen served as a desolvation and a cone gas with flow rates of 800 L ⋅ h^−1^ at 350 °C and 25 L ⋅ h^−1^. The mass spectrometer operated in the multiple reaction monitoring mode using argon as a collision gas at a pressure of ∼3 x 10^−3^ bar. Cone voltage and collision energy were optimized for maximum signal intensity of each individual compound during ionization and collision-induced dissociation with a dwell time of 0.025s per transition. In some samples, (Fig. 2d-e), sucrose and glucose were measured enzymatically in chloroform-methanol extracts.

### Amino acid measurements

Around 50 mg (fresh weight) of shoot or root material of individual genotypes were frozen in liquid nitrogen and ground to powder. To each sample, 300 µl of extraction solvent (50% (v/v) MeOH, 1.07 mg/ml methyl methanethiosulfonate, 10 µM norvaline, 50 mM HCl) was added. Samples were shaken for 10 min at 21 Hz in a cooled mixer mill. After centrifugation at 4°C and 18,000 x *g* for 10 min, 300 µl of the supernatant were transferred to a fresh tube. The pellet was then reextracted with 300 µl extraction solvent as before and the supernatants combined. After centrifugation (4°C and 18,000 x *g* for 10 min), 20 µl of the final supernatant were then derivatized using the AccQ-Tag Ultra Derivatization Kit (Waters Corporation) according to the manufacturer’s instructions. An aliquot (50 µl) of the derivatized amino acid extract was transferred to UPLC vials. AAs were measured as described previously^52^.

### Fluorometric starch quantification

Total starch content was measured from rosette leaves of 5-weeks-old *A. thaliana* WT, and bZIP triple (*bzip2/-11/-44*) or quintuple (*bzip1/-2/-11/-44/-53*) mutants grown under long-day conditions at 100 µmol photons m^-2^ s^-1^ and 60% relative humidity using an Amplex Red-based fluorescent quantification assay. To this end, 50 mg of fresh plant material were homogenised and extracted three times with 1 ml of 90% (v/v) ethanol and incubation for 5 min at 60°C to remove soluble glucose and other small oligosaccharides. After centrifugation at 18,000 x *g* for 10 min, the supernatant was discarded, and the remaining starch-containing pellets were resuspended in 500 µl of 0.5 M KOH and incubated at 100 °C for 5 minutes. Subsequently, the pH was adjusted to 4.8 by adding 350 µl of 0.4 M H_3_PO_4_. Following the addition of 100 µl of an enzyme mix containing 0.5 U α-amylase (Sigma-Aldrich, Catalogue-No. A3176 https://www.sigmaaldrich.com) and 7 U amyloglucosidase (Sigma-Aldrich, Catalogue-No. 10115) dissolved in 50 mM phosphate buffer (pH 7.4), samples were incubated for one hour at room temperature to allow enzymatic starch hydrolysis. Following centrifugation as mentioned above, the supernatants containing the resultant glucose moieties were transferred to fresh tubes and aliquots thereof used for the subsequent fluorometric glucose quantification. The reaction mixture consisted of 0.5 µl 10 mM Amplex Red (Invitrogen, Catalogue-No. A22188; https://www.thermofisher.com), 1 µl horseradish peroxidase (10 U/mL, Sigma-Aldrich, Catalogue-No. 77332), 1 µl glucose oxidase (100 U/mL, Sigma-Aldrich, Catalogue-No. G7141), 47.5 µl 100 mM phosphate buffer (pH 7.4) and 50 µl of a given sample (diluted 1:40 or 1:60 in 100 mM phosphate buffer, pH 7.4). Fluorescence was detected after a 30-minute incubation at room temperature in the dark using a Fluoroskan Ascent microplate reader (Thermo Fisher Scientific, Waltham, USA) at an excitation wavelength of 530 nm and an emission wavelength of 590 nm. Quantification was based on a calibration curve obtained from freshly hydrolysed pure starch (Sigma-Aldrich, Catalogue-No. 85642) used at four different concentrations.

### RNA-sequencing

To obtain a global transcript profile of WT, *bzip2/-11/-44* and *bzip1/-2/-11/-44/-53*, we cultivated plants at 100 µmol m^-2^ s^-1^ under long-day conditions until the onset of flowering of the primary inflorescence. At this developmental stage, we harvested at ZT0, flowers, mature and young leaves as well as an axillary branch enriched tissue (rosette core) from three individual plants per genotype and pooled the material to create each biological replicate. Total RNA was extracted from three biological replicates per genotype and tissue using the Trizol extraction method^53^. RNA quality was checked using a 2100 Bioanalyzer with the RNA 6000 Nano kit (Agilent Technologies; https://www.agilent.com). The RNA integrity number for all samples was g7.7. DNA libraries suitable for sequencing were prepared from 500 ng of total RNA with oligo-dT capture beads for poly-A-mRNA enrichment using the TruSeq Stranded mRNA Library Preparation Kit (Illumina; https://www.illumina.com) according to manufacturer’s instructions (½ volume). After 13 cycles of PCR amplification, the size distribution of the barcoded DNA libraries was estimated to be approximately 310 bp by electrophoresis on Agilent DNA 1000 Bioanalyzer microfluidic chips. Sequencing of pooled libraries, spiked with 1% PhiX control library, was performed at ∼25 million reads/sample in single-end mode with 100 nt read length on the NextSeq 2000 platform (Illumina) using a P3 sequencing kit. Demultiplexed FASTQ files were generated with bcl2fastq2 v2.20.0.422 (Illumina). Quality control of the sequencing data was performed by assessing the FastQC report. Reads of each sample were mapped to the Arabidopsis genome (Col-0 TAIR10 genome release) as reference using salmon v0.7.2^54^. In all samples, between 70 and 85 % of the reads could be unambiguously mapped. Only mapped reads were used to calculate the relative abundance in transcripts per million (TPM) units. TPM values were used for downstream analyses with R software (www.cran.r-project.org, R version 3.5.1) and the GenomicRanges, rtracklayer^55^, samtools 0.1.18, and edgeR^56^ packages. Differences in gene expression between WT and mutants were separately analyzed for each WT/mutant comparison using the DESeq2 package^57^. Genes with an adjusted p-value f 0.05 (method: p-adjust <BH= correction)^58^ and an absolute log_2_FC f –1 (down-regulated) or log_2_FC g 1 (up-regulated) were considered as differentially expressed (Supplementary Table 2). Raw data from RNAseq of this article can be found in Gene Expression Omnibus (GSEnumber?).

### Quantitative realtime PCR

For transcript analysis total RNA was extracted from plant tissue using the Trizol extraction method^53^. RNA was converted to complementary DNA using H^-^ reverse transcriptase and a mix of random nonamer and oligo dT primers. Quantification of transcript abundance was done by quantitative realtime PCR (qRT-PCR) using SYBR green-based detection as described before^59^. Data were obtained from three biological replicates each derived from plant material of three individual plants. Presented are mean expression values normalised to *UBIQUITIN5* expression. Gene specific oligonucleotide primers used for qRT-PCR analyses are summarised in Supplementary Table 3.

### ATAC-sequencing

Four-week old WT and *Pro35S*::*bZIP11*-*GR* plants were used to extract leaf protoplasts via the epidermal leaf peel method^60^. Six leaves were placed in 10 ml of enzyme solution (1 % (w/v) cellulase ‘Onozuka’ R10, 0.25 % maceroenzyme ‘Onozuka’ R10, 0.4 M mannitol, 20 mM KCl, 20 mM MES-HCl, 10 mM CaCl_2_, 0.1 % (w/v) BSA, adjusted to pH 5.7) and digested for 1 h at room temperature with constant gentle agitation. Cells were filtered through a 50 μm mesh and washed twice in W5 solution (154 mM NaCl, 125 mM CaCl_2_, 5 mM KCl, 2 mM MES-HCl adjusted to pH 5.7). Protoplasts were re-suspended in MMg (0.4 M mannitol, 15 mM MgCl_2_, 4 mM MES-HCl, adjusted to pH 5.7) at a concentration of 200,000 cells per ml. Reactions took place in 2 ml in six-well plates with constant agitation. WT and *Pro35S*::*bZIP11*-*GR* cells were treated with 10 μM DEX and mock with solvent (acetone). After treatment for 45 min, cells were collected by centrifugation for ATAC-seq library preparation. ATAC-seq library preparation was performed as described, previously^61, 62^. Following protoplast extraction, nuclei were isolated from approximately 50,000 cells per reaction by sucrose sedimentation^63^. Freshly extracted cells were centrifuged at 500 x *g* at 4°C for 10 min. The following steps were all carried out on ice. The supernatant was discarded, and the pellet was resuspended in 1 ml of ice-cold nuclei purification buffer (20 mM MOPS-HCl, 40 mM NaCl, 90 mM KCl, 2 mM EDTA, 0.5 mM EGTA, 0.5 mM spermidine, 0.2 mM spermine, 1 x protease inhibitors, adjusted to pH 7). Cells were then filtered through a 30 μm mesh. Nuclei were collected by centrifugation at 1200 x *g* for 10 min at 4°C and the pellet resuspended in 1 ml of ice-cold nuclei extraction buffer 2 (0.25 M sucrose, 10 mM Tris-HCl, pH 8, 10 mM MgCl_2_, 1 % (v/v) Triton X-100, 1 x protease inhibitors). This step was repeated however, this time the pellet was resuspended in 300 μl of NPB and the resuspended nuclei were carefully layered over 300 μl of ice-cold nuclei extraction buffer 3 (1.7 M sucrose, 10 mM Tris-HCl, pH 8, 2 mM MgCl_2_, 0.15 % (v/v) Triton X-100, 1 x protease inhibitor). The two layers were then centrifuged at 300 x *g* for 20 min at 4°C and the supernatant was removed. Nuclei were resuspended in 50 μl of tagmentation reaction mix as per manufacturer instructions (TDE1, Illumina) and incubated at 37°C for 30 mins with gentle agitation every 5 min. Reactions were purified following the manufacturer’s instructions using a QIAGEN MiniElute PCR purification kit (https://www.qiagen.com) and eluted in 11 μl of elution buffer. DNA was amplified by PCR using ATAC barcoded primers and NEB Next High Fidelity PCR Master Mix (5 min 72°C, 30 sec 98°C, then 5 x (10 sec 98°C, 30 sec 63°C, 1 min 72°C) held at 4°C). An aliquot (5 μl) of the PCR reaction was then further amplified by qPCR (30 sec 98°C, then 20 x (10 sec 98°C, 30 sec 63°C, 1 min 72°C)) to determine the required number of additional cycles. Additional cycle number for each reaction was determined by the cycle number for which a reaction has reached one third of its maximum, using the linear fluorescence vs. cycle number graph from the qPCR. All libraries were purified with AMPure XP beads at a ratio of 1.5: 1 beads: PCR reaction. Libraries were eluted in 20 μl of 10 mM Tris, pH 8. Libraries were sequenced using the Illumina HiSeq paired end 150 bp method by NovoGene, Singapore (https://www.novogene.com). Processing was carried out using Galaxy Australia^64^ and R with RStudio (Version 4.2.2) with the following steps. In Galaxy, raw reads were trimmed using Trimmomatic^65^ with a 10 bp HEADCROP, a SLIDINGWINDOW with an average quality of 30 over every 6 bp, and an ILLUMINACLIP NexteraPE. Reads were mapped against the *Arabidopsis thaliana* TAIR10 reference genome using Bowtie2^66^, with paired end, dovetailing, and a maximum fragment length of 1000. Reads smaller than 30 bp, duplicate reads, reads with a quality score of <30 phred, and those which were mapped to the chloroplast or mitochondrial genome were discarded. Peaks were called with MACS2^67^, using the inputs: single-end BED, effective genome size 1.2^e8^, an extension size of 200 and a shift size of 100. BED, BAM and index files were then imported into RStudio and DARs were determined using the package DiffBind. Peaks were read with peakCaller=“narrow”, minOverlap=3 and dba.contrast function was specified to compare mock treated and bZIP11-induced samples. The package rtracklayer was used to convert the DiffBind peaks report into BED format. The peaks report was then imported back into Galaxy where differential peaks were annotated to the *Arabidopsis* TAIR10 reference genome using ChIPseeker^68^. The resulting BED file of annotated DARs was imported into the Interactive Genome Viewer (Broad Institute, University of California)^69^ along with BED files of samples from MACS2 output for visualisation.

### DAP-sequencing

DAP sequencing data were obtained online from the Anno-J Networked Genome Browser (http://neomorph.salk.edu/dap_family.php?active=DAP+data/bZIP_tnt). A detailed description of the DAP-Seq experiment was published previously^35^.

### Protoplast Transactivation assays

Protoplast transfection assays were performed as previously described^70^ with minor modifications^21^. β-glucuronidase (GUS) enzyme assays were performed after 16 h incubation. Construction of effector plasmids was described previously^23^. *SWEET15* and *GAT1_2.1* reporter constructs were generated in this study, by PCR-amplifying 1.6 kb of the *SWEET15* and 1 kb of the *GAT1_2.1* promoter using sequence specific primers that introduced NcoI and PstI restriction sites into the PCR product. After restriction of the PCR products with NcoI and PstI, the resulting promoter fragments were ligated into the multiple cloning site of the pBT10-GUS vector, upstream of the *GUS* reporter gene. For mutagenesis of putative bZIP binding sites within the *SWEET15* and *GAT1_2.1* reporter constructs, the QuickChange II site-directed mutagenesis kit (Agilent Technologies) was used, in combination with the mutagenic primers given in Supplementary Table 3.

### Statistics

Statistically significant differences between genotypes at individual timepoints were assessed by two-tailed Student’s *t*-test (n.s., not significant; *pf0.05, **pf0.01, ***pf0.001) and are presented either in the figures or for reasons of clarity given in Supplementary Table 4.

## Acknowledgements

We thank Prof. Dr. Wolfgang Droege-Laser (University of Wuerzburg, Germany) for constant support and proof reading, Theresa Damm (University of Wuerzburg, Germany) for excellent technical assistance and the Core Unit SysMed at the University of Wuerzburg for excellent technical support and RNA-seq data generation. We thank Prof. Dr. George Coupland (MPI for Plant Breeding Research, Cologne) for providing transgenic lines. This work was supported by the IZKF at the University of Wuerzburg (project Z-6). PK was supported by a research grant from the German Research foundation to CW (DFG project 408153945). We thank the Wuerzburg graduate school of life sciences (GSLS) for constant support and funding via the PostDoc plus program to CW. CAB was funded by an Australian Research Council (ARC) Georgina Sweet Laureate Fellowship (FL180100139) and the ARC Centre of Excellence for Plant Success in Nature and Agriculture (CE200100015). AMH was supported by a Commonwealth Scientific and Industrial Research Organization Postgraduate Scholarship. RF and JEL were supported by the Max Planck Society.

---

**Fig. S1.**
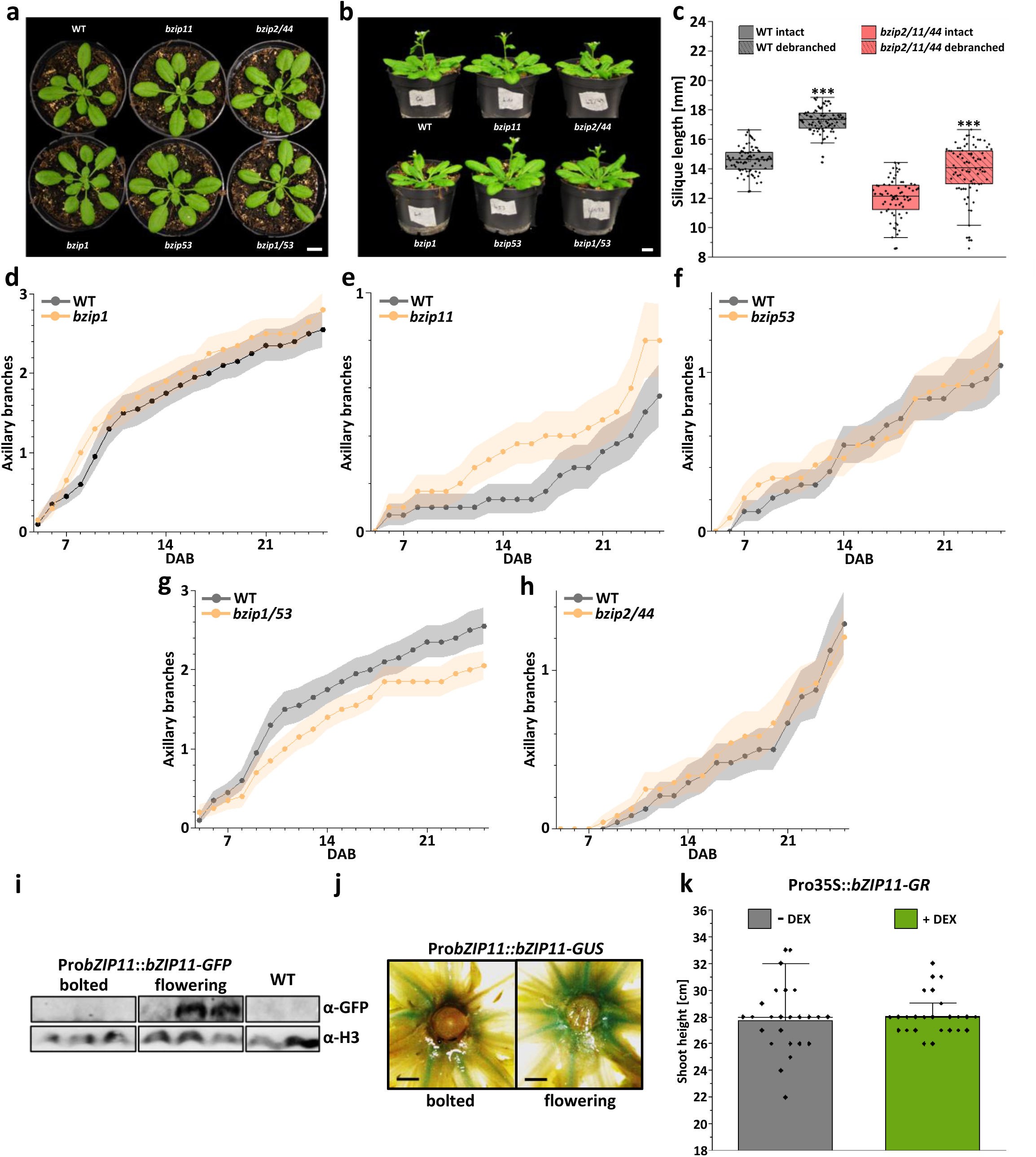
I Growth phenotypes of single and double mutant combinations of S_1_-bZIP TFs and bZIP11 expression kinetics. Vegetative and reproductive growth of single and double mutants of S_1_-bZIP TFs. Single (*bZIP1*, *bZIP11*, *bZIP53*) or double mutants (*bZIP2/-44*, *bZIP1/-53*) of group S_1_ bZIP TFs do not show obvious differences in (**a**) vegetative or (**b**) reproductive growth. Given are representative pictures of (**a**) 4– to (**b**) 5-week-old plants. Scale bar: 1cm. **c**, Silique length (n=70-80 per genotype and treatment) of siliques 6 to 15 (from the inflorescence base) from 12-week-old WT (black box) or *bzip2/-11/-44* mutant plants (red box) that were either kept untreated (intact, clear boxes) or from which all cauline or axillary branches were removed upon outgrowth initiation (debranched, striped boxes). **d-h,** Mean axillary branching (± SEM, n=20-30 per genotype) of WT (black line) or single (**d**) *bZIP1*, (**e**) *bZIP11*, (**f**) *bZIP53,* and double (**g**) *bZIP1/-53*, (**h**) *bZIP2/-44* mutants of S_1_ bZIP TFs (yellow line). Outgrown axillary branches (≥1cm) were counted starting 5 days after bolting (DAB). **i,** bZIP11 protein abundance in rosette cores of just bolted or flowering WT and Pro*bZIP11*::*bZIP11*-GFP plants, respectively, was determined by Western-Blot analysis using a GFP specific antibody (α-GFP). Histone H3 detection (α-H3) was used to demonstrate equal loading. **j**, Expression of bZIP11 in rosette cores and leaf petioles at ZT8 was demonstrated by histological GUS staining using Pro*bZIP11*:*bZIP11*-GUS plants. Given are representative pictures from 3 repetitions. Scale bar: 1mm. **k**, Local overexpression of bZIP11 in rosette cores does not affect primary shoot growth. Given is the mean shoot height (± SEM, n=24) of recurringly mock treated (– DEX, grey bar) or DEX induced (+ DEX, green bar) Pro*bZIP11*::*bZIP11*-GR plants at 16 days after start of induction (DAI). Plants were grown on soil (**Fig. S1a-k**) under long day conditions at 100 µmol photons m^-2^ s^-1^. Statistically significant differences were determined by Student’s *t*-test comparing intact and debranched plants (**Fig. S1c**) or WT and respective bZIP mutants (for **Fig. S1d-h**, **k**, see **Supplementary Table 4**), *** P<0.001.

**Fig. S2.**
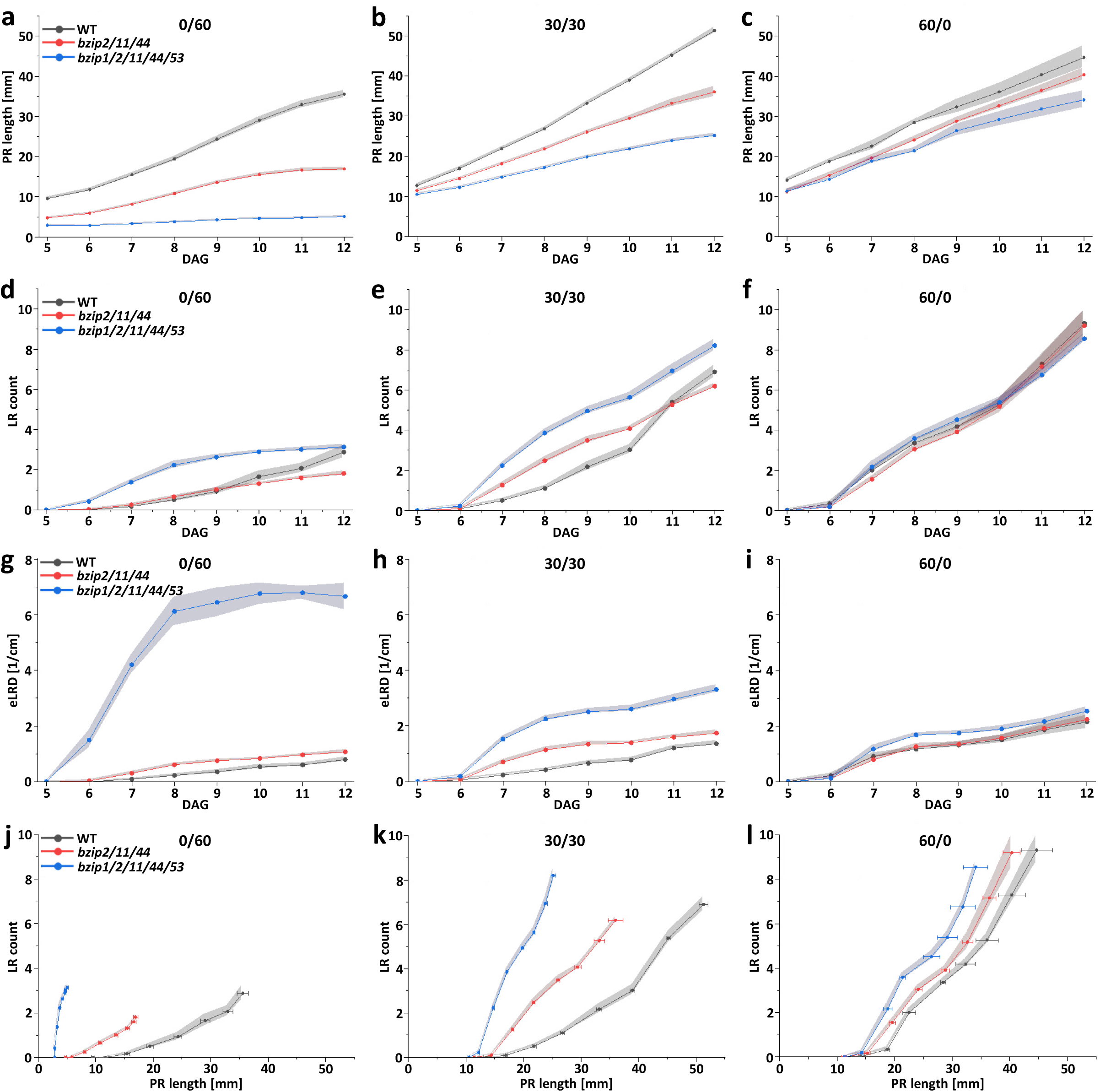
I Sucrose feeding largely rescues root phenotypes of S_1_ bZIP mutants. **a-l**, Root architecture parameters of WT (black line) bZIP triple (red line) and quintuple (blue line) mutants grown for up to 12 days after germination (DAG) under long day conditions on ½ strength MS medium supplemented with either 60 mM mannitol (0/60), 30mM sucrose + 30 mM mannitol (30/30) or 60 mM sucrose (60/0). The poorly metabolised mannitol serves as osmotic control. **a-c**, Primary root (PR) length, **d-f**, Lateral root (LR) number, **g-I**, Emerged lateral root density (eLRD), **j-l**, Emergence of LRs during PR growth. Given are mean values (± SEM, n=6, each replicate consisting of 10 plants per genotype and treatment). Statistically significant differences were determined by Student’s *t*-test comparing WT and respective bZIP mutants at (**a-i**) specific timepoints or (**j-l**) PR length (see **Supplementary Table 4**).

**Fig. S3.**
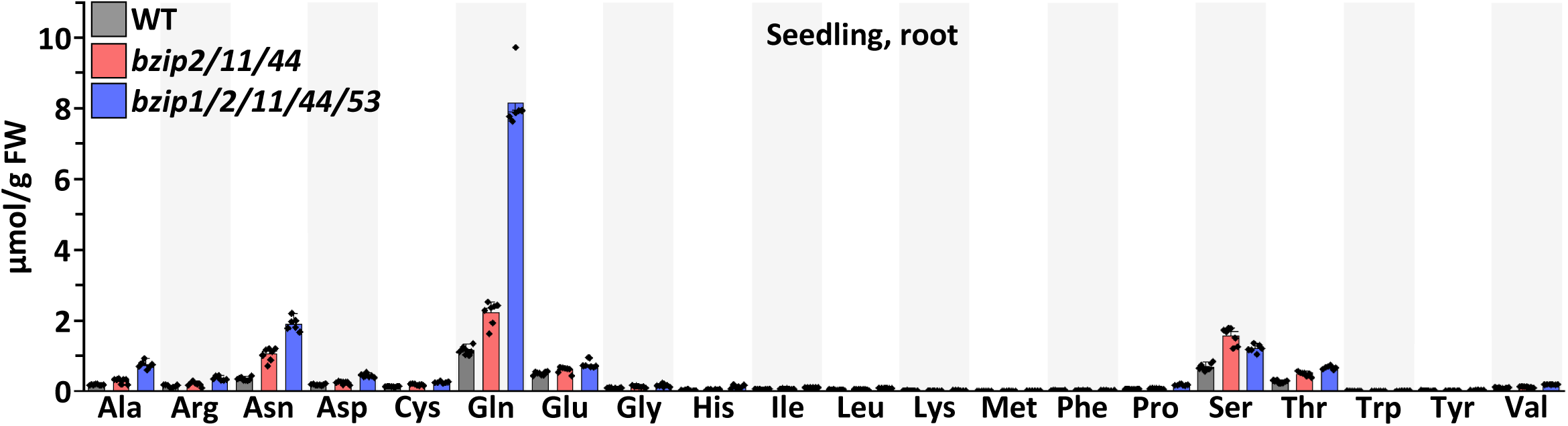
I S_1_ bZIP mutants show a marked increase in glutamine levels in roots. Given is the mean amino acid content (bar box plots, n=6-8) determined by UPLC-MS/MS in roots of 3-week-old WT (black box), bZIP triple (red box) and quintuple mutant (blue box) seedlings. Root material of several plants was harvested at ZT8 and pooled. Plants were grown on ½ strength MS medium under 12h light / 12h dark conditions at 100 µmol photons m^-2^ s^-1^. Statistically significant differences were determined by Student’s *t*-test comparing individual amino acid content of WT and respective bZIP mutants (see **Supplementary Table 4**).

**Fig. S4.**
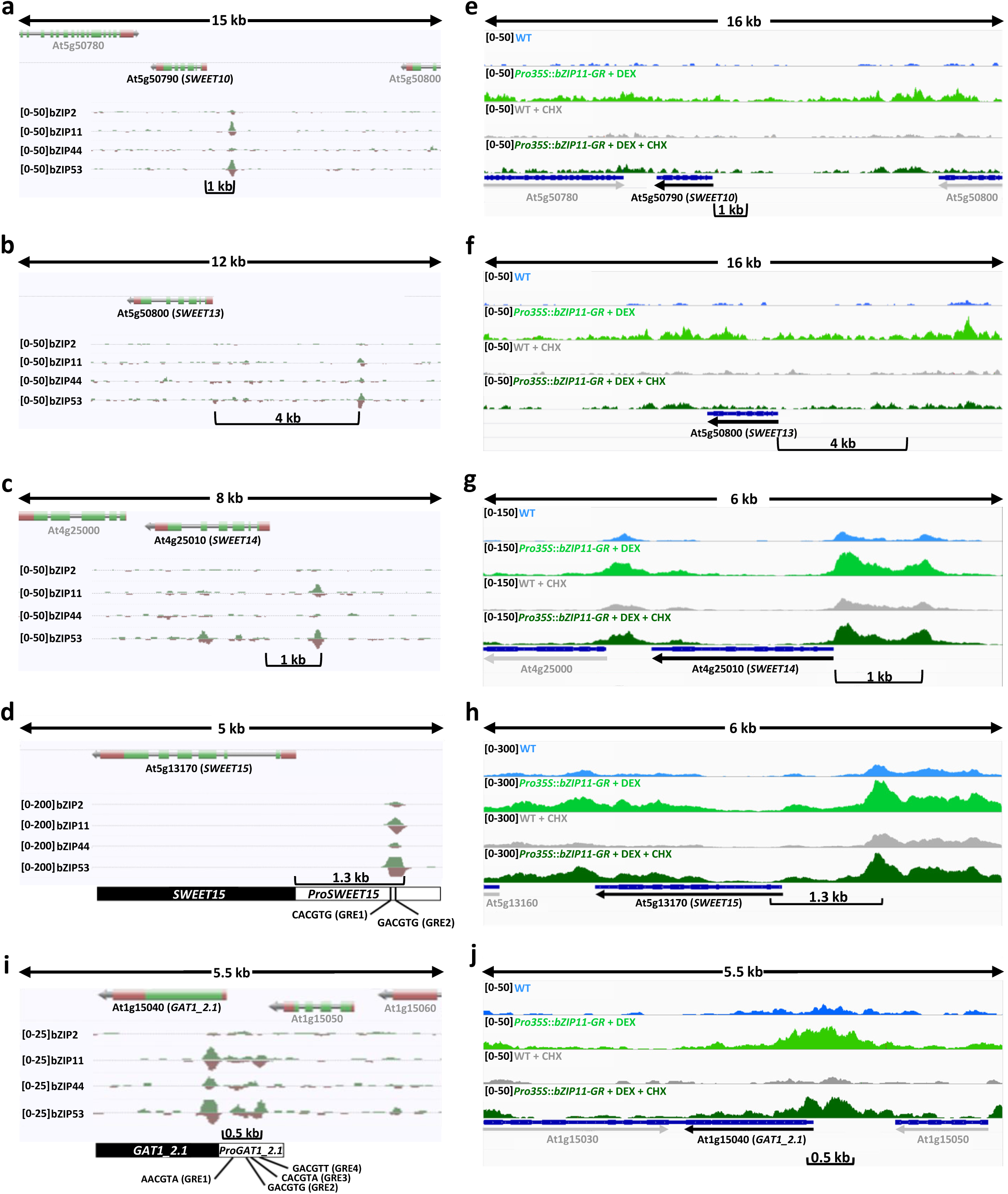
I Binding of S_1_ bZIP TFs to target promoters and their impact on promoter chromatin accessibility. **a-d**,**i**, DNA affinity purification sequencing (DAP-seq) of S_1_-bZIPs (bZIP2, –11,-44,-53) (O’Malley et al., 2016) reveals direct bZIP binding to promoters of specific clade 3 *SWEETs* (**a**, *SWEET10*; **b**, *SWEET13*; **c**, *SWEET14*; **d**, *SWEET15*) or (**i**) *GAT1_2.1*. Position of typical bZIP binding sites (G-box related elements (GREs) are given. **e-h**, **j**, Assay for Transposase-Accessible Chromatin using sequencing (ATAC-seq) from protoplasts derived from WT and dexamethasone induced (+ DEX) *Pro35S*::*bZIP11*-GR plants in absence or presence of cycloheximide (+ CHX) reveals bZIP11 dependent chromatin opening in promoters of specific clade 3 *SWEETs* (**e**, *SWEET10*; **f**, *SWEET13*; **g**, *SWEET14*; **h**, *SWEET15*) or (**j**) *GAT1_2.1*.

**Fig. S5.**
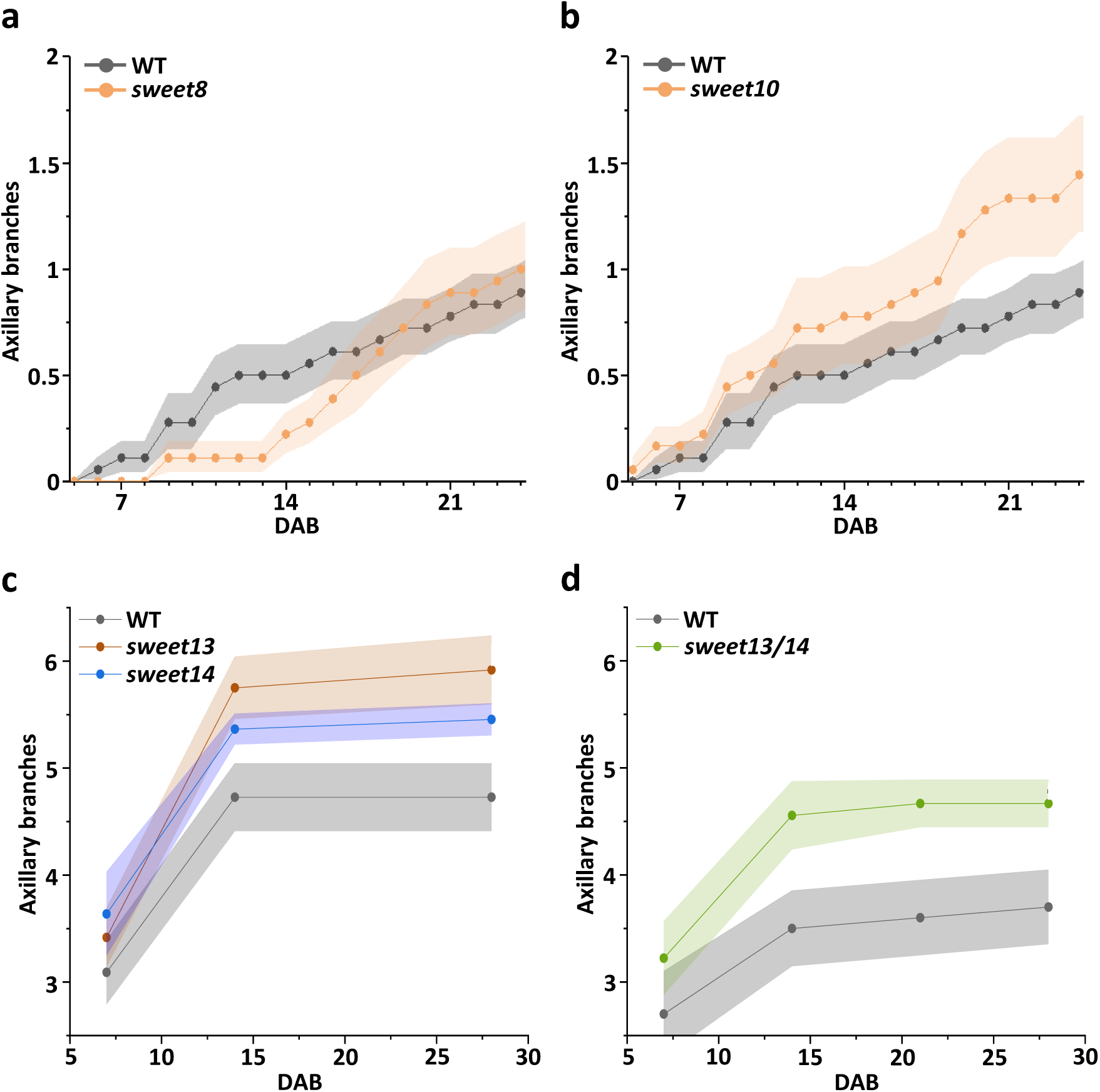
I Axillary branching phenotypes of sweet*s, –10, –13, –14* single and *sweet13/-14* double mutants. Mean axillary branching (± SEM, **a,b**, n=18 or **c,d** n=10 per genotype) of WT (black line) and single (**a**) *sweet8* (yellow line), (**b**) *sweet10* (yellow line), (**c**) *sweet13* (brown line) and *-14* (purple line) and (**d**) double *sweet13/-14* plants (green line), respectively. Plants were grown on soil under long day conditions at 100 µmol photons m^-2^ s^-1^. Outgrown axillary branches (≥1cm) were counted starting 5 to 7 days after bolting (DAB). Statistically significant differences were determined by Student’s *t*-test comparing WT and respective mutants (see **Supplementary Table 4)**.

